# Discovery of potential epigenetic inhibitors against histone methyltransferases through molecular docking and molecular dynamics simulations

**DOI:** 10.1101/2020.04.13.040030

**Authors:** Tirumalasetty Muni Chandra Babu, Zaiping Zhang, Danian Qin, Chengyang Huang

**Author notes:** Equal contribution, T.M.C.B. and Z.P.Z. contributed equally in this work. Corresponding author: Danian Qin and Chengyang Huang.

## Abstract

Histone methyltransferases (HMTases) catalyze histone methylations that are the important epigenetic marks regulating gene expression, cell fate, and disease progression. In this study, we investigated potential epigenetic inhibitors against HMTases through in silico approaches, including ensembled molecular docking and molecular dynamics simulations (MDS).We identified three candidate compounds, including echinomycin, emetine, and streptonigrin, which showed interactions with HMTases. Echinomycin showed similar binding affinity with H3K4-HMTase NSD3 and H3K9-HMTase SETDB1 but streptonigrin and emetine had preferential binding affinity with NSD3 and SETDB1, respectively. The binding of NSD3 to streptonigrin and echinomycin and binding of SETDB1 to emetine and echinomycin were further confirmed by the results of hydrogen bonding profile and MM/PBSA calculations. Together, our results uncover the binding affinities of echinomycin, emetine, and streptonigrin with histone methyltransferases, and suggest that these compounds are potential epigenetic inhibitors regulating cell activities.

## 1. Introduction

Chromatin is a highly condensed compact structure responsible for all genetic processes inside the nuclei of eukaryotic cells [1]. The basic unit of chromatin is a nucleosome composed of ~147 base pairs of DNA wrapped around an octameric core of histone H2A, H2B, H3, and H4 [2, 3]. It is well established that the flexible N-terminal tails of histones from the nucleosome are the targets of the different posttranslational processes such as methylations of lysine and arginine, phosphorylation of serine, threonine ligation of protein domain and ubiquitination [4]. Histone methylations of H3K4, H3K36, and H3K79 are associated with open chromatin and are involved in gene activation, whereas methylations of H3K9, H3K27 and H4K20 are predominant in heterochromatin formation and may repress the gene expression [5, 6].

Investigation on molecular signaling pathways of cancer and stem cells has led to the discovery of novel target-based therapeutic strategy targeting epigenetic regulators. In recent years, many of the advanced structural genomic and proteomic analysis has led to exploration of new therapeutic target histone methyltransferases (HMTase) that regulate gene expression by the epigenetic alterations [7,8]. Histone methyltransferases (HMTases) are histone-modifying enzymes, which transfer of one, two or three methyl groups from its cofactor, S-adenosyl-L-methionine (SAM) to lysine or arginine side chain of histone, H3 protein. Subsequently, changes in histone methylations regulate chromatin structures and components of chromatin-binding protein complexes that influence gene transcription and downstream signaling. Aberrant histone methylations are relevant events in many types of cancers [9].

HMTases involved in H3K4 and H3K9 methylations have been associated with cancer, genetic disorders and many life-threaten diseases with the deregulation of their activities. HMTase like MLL3, MLL4 and NSD3 are involved in hepatocellular carcinoma [10], colorectal cancer [11], and acute myeloid leukemia [12]. While GLP has been found to cause medulloblastoma by its downregulation [13] and 9q sub-telomeric deletion syndrome [14]. Hepatocellular carcinoma and Huntington’s disease are caused by the overexpression of SMYD2 and SETDB1, respectively [15, 16]. However, the disease mechanism and development were not well established yet [17].

Methylation of lysine or arginine residue is a biological hallmark found in the conserved catalytic domain of SET (Suppressor of variegation, Enhancer of Zesta trithorax) which is composed of ~130 amino acids. HMTase has two distant sites on a surface of the SET domain for the binding of methyl donating cofactor and the methyl-accepting substrate, respectively [18]. The substrate binding site has enclosed with the deep cavity and is structurally druggable and diverse since different substrates have been recognized by different HMTases. Moreover, HMTase substrate competitive inhibitors can also be selective. For example, UNC0638 inhibits G9a and GLP but not SUV39H1 and SUV39H2 [19]. All HMTases share the same pocket of the cofactor binding site for SAM, although they are structurally diverse and comparable with the ATP-binding pocket of kinases [20]. These are the two vital binding sites in HMTase that have become a challenge for drug discovery and development of target specific inhibitors.

To identify the potent HMTases inhibitors, we focus on bioactive compounds derived from natural sources. Natural products and their derivatives are structurally complex because they cover much larger space and have many chiral centers and functional groups that allow them to achieve higher bioactivity [21]. In this scenario, we used in-house compound library of 502 natural compounds in the current study due to their wide range of pharmacological properties in cancer, neurodegenerative diseases, and aging process. Earlier studies have been reported that some of the natural compounds are found to inhibits HMTase few examples are as follows. Protoberberine, an alkaloid and gliotoxin analogs from marine fungus have shown the inhibitory activity of HMTase, G9a [22, 23] However, the inhibition of many other HMTases by the natural compounds are still unknown.

Because of the demand for natural product-based HMTase inhibitors and the extreme importance of future epigenetic therapies, there is a clear need to screen novel scaffolds from the natural sources that could be served as potential leads by regulating the histone methylation. Our study majorly aimed to investigate epigenetic compounds against HMTases by analyzing their molecular interactions with various HMTases.

## 2. Materials and Methods

### 2.1. Chemicals and reagents

Echinomycin (Cayman Chemicals Ann Arbor, MI, USA), Emetine, Methylcaconitin, Streptonigrin, Brefeldin A, Harringtonine and Dehydroandrographolide (abcam, Cambridge, UK), Dimethylsulfoxide (DMSO) (MP Biomedicals, lllkirch, France), MTT (3-[4, 5- dimethyl thiazol-2-yl]-2,5 diphenyl tetrazolium bromide) (Alfa Aesar Chemicals Co. Ltd. Shanghai, China), fetal bovine serum (FBS) (Hyclone Laboratories, Logan, UT, USA), DMEM (Dulbeccos modified eagles media) (Hyclone Laboratories, Logan, UT, USA), specific antibody for H3K4me3, H3K9me3 (abcam, Cambridge, UK) and GAPDH (ZSGB-BIO, Beijing, China).

### 2.2. Culture of glioma cells

Human glioma cells U251 cell line was procured from the ATCC (American Type Culture Collection) and cultured as a monolayer at 60-70 % of confluence in the low glucose level Dulbecco’s modified eagles media with the supplements of 10 % fetal bovine serum and 1 % penicillin-streptomycin were used for further sub-cultures. The cells were incubated in a 95 % air and 5 % of CO2 humidified atmosphere at 37°C.

### 2.3. Cytotoxic assay

The cytotoxic ability of 502 natural compounds of in-house library was evaluated on U251 glioma cells through broad range cytotoxic assay. Initially, the cells were seeded in the 96-well microtiter plate at 6×103 cells per well and incubated the cultures at 37°C for overnight to obtain the required confluence. After the incubation, all the compounds were added individually in the final concentration of 20 µg/mL and further incubated for 48 h. Subsequently, the inhibition of cell viability was quantified by MTT assay. The MTT reagent in the concentration of 0.5 mg/mL was dissolved in DMSO and added into each well, followed by incubation at 37°C for 4 h in dark conditions. Further, the viable cells were detected by reading the absorbance of tetrazolium product, formazan at 570 nm using the TECAN Infinity 200pro microplate reader. The viability of cells and cell count were compared with negative control (treated with DMSO) and analyzed quantitatively [24].

### 2.4. Western blotting

The best hit compounds with proven cytotoxic effect were used to quantify their effect on the tri-methylation of histone H3 viz. H3K4me3 and H3K9me3 through the western blotting as per the regular protocols.

The overnight incubated cells as processed in the cytotoxic assay were taken for the treatment of compounds. The cells were treated the aforementioned seven compounds individually in the concentration of 20 µg/mL and incubated for 48 h. After the incubation, the cells were subjected to lysis with SDS-loading buffer (50 mM Tris-HCl pH 6.8; 2% w/v SDS; 10% glycerol) and boiled for 10 min. The protein samples were analyzed with 15 % sodium dodecyl sulfate- polyacrylamide gel electrophoresis (SDS-PAGE) and transferred to the nitrocellulose membrane. The membrane was blocked with 5% w/v nonfat dry milk for 1 h at room temperature subsequently probed with mouse monoclonal antibody specific for H3K4me3 and H3K9me3 (1: 2000; abcam) and rabbit polyclonal antibody specific for GAPDH (1:1000; ZSGB-BIO) and incubate for 16 h at 4°C. Further, the membrane was washed with phosphate buffer and again incubated with 680 red fluorescence and 800 green fluorescence-conjugated secondary antibodies (1:7000; ZSGB-BIO) for 1 hr. The expressed protein was visualized through an Odyssey Infrared Imaging system (LI-COR, Biosciences, Lincoln, USA).

### 2.5. Molecular docking analysis of HMTases with selective compounds

In order to enumerate the best lead molecule, docking analysis was performed on these selected seven compounds, emetine (EMT), methylcaconitin (MLA), streptonigrin (SYN), brefeldin A (BFA), harringtonine (HRT) and echinomycin (ECN) and dehydroandrographolide (DHA) with epigenetic regulatory enzymes of H3K4 methylation (SETMAR, SETD7, NSD3, SMYD2, MLL1, MLL3, MLL4, and MLL5); H3K9 methylation (GLP, G9a, SUV39H1, SUV39H2, and SETDB1).

The experimental procedure was carried out using the docking module implemented in MOE 2010.12 (Molecular Operating Environment) [26]. All the protein co-crystal structures were retrieved from the protein data bank (www.rcsb.org). Initially, protein preparation was carried out by the addition of polar hydrogens to protein structure with the protonation process. To obtain the stable conformer of the protein structure, energy minimization was performed using the MMFF94x force field. The inhibitor binding site residues were collected from literature, which is selected and highlighted through “Site Finder” module implemented in MOE software, and docking was carried out with default parameters, that is, placement: triangle matcher, Recording 1: London dG, Refinement: Force field and a maximum of 10 conformations of each compound was allowed to save in a separate database file in. mdb format.

The binding energy and binding affinity of protein-ligand complexes were calculated using molecular mechanics generalized Born interactions/volume integral (MM-GB/VI) implicit solvent method in MOE [27]. Non-bonded interaction energies between the receptor protein and ligand molecules include Van der Waals, Coulomb, and Generalized Born implicit solvent interactions energies are categorized as Born interaction energy. The binding energy and affinity values were calculated for each compound against various target proteins and reported in the unit of kcal/mol.

The Protein-protein docking was also employed to H3K4 and H3K9 peptide with selected HMTase using the HDOCK server [28] to reveal the dynamic behavior of protein and peptide complexes when compared to native protein and compound complexes.

### 2.6. Molecular dynamics (MD) simulations

Molecular dynamics simulations (MDS) were performed using GROMACS 5.1.2 [29] for 50 ns time period with the OPLSA force field. To enumerate the thermodynamic behavior of histone methyltransferase and elucidate the mechanism of its inhibition, MDS was carried out in the combinations of protein alone and protein complexes with ligand, H3K4 peptide and H3K9 peptide. The topology of protein was created using the OPLSA program and ligand topology parameterization was created using LigParGen [30] an OPLS-AA parameterization server for organic ligands.

The cubic water box was created with a simple point charge water model [31] with the appropriated dimensions based on the surface area of the protein structure. The protein structure (liganded and unliganded protein) was immersed in the center of the cubic box with a minimum distance of 1.0 nm between the wall of the water box and the surface of any part of the protein was set up for the initial process of solvation in the MD run. Further, the solvated system was neutralized with an ionic strength of 0.1 M, Na+ (sodium) and Chloride (Cl-) ions with replacing appropriated water molecules. The neutralized system was minimized through the algorithm steepest descent, minimization with a maximum number of steps 50000, long-range of electrostatic interactions was computed by particle mesh Ewald (PME). The short-range electrostatic and van der walls cut-off was set with 1.0 nm. The system was equilibrated under NVT conditions (constant number, volume, and temperature) for 500 ps at 300 K under isothermal ensemble by soft coupling with Berendsen thermostat [32]. The internal time step was 2 femtosecond (fs) and update the log file of every 50.0 ps. While in NPT conditions (constant number, pressure, and temperature) periodic boundary conditions were used with a constant number of particles in the systems constant pressure and the constant temperature criteria. In this simulation, the system was coupled with Parrinello-Rahmanbarostat equilibrate at 1 bar pressure for 500 ps [33]. The production dynamics for 50 ns was performed for protein and protein complexes with the compounds, H3K4 and H3K9 peptides and trajectories were sampled at every 50 ps interval.

The analysis of MD trajectories such as RMSD, Rg, RMSF and SASA was carried out using GROMACS tools. The protein secondary structure analysis and visualization was carried out using VMD timeline-plugin implemented in visual molecular dynamics (VMD) [34].

### 2.7. Binding free energy calculation by MM/PBSA

The protein-ligand binding free energy and energy contribution of each amino acid residue of HMTase were computed using GROMACS plug-in g_mmpbsa program through MM/PBSA calculations [35]. The binding free energy of protein-ligand complex in the solvent is calculated as G_bind_ = G_complex_-[G_protein_ + G_ligand_]

### 2.8. Cloning, Expression, and purification of human SETBD1

The synthetic gene encoding catalytic domain of human histone methyltransferase, SETDB1 (196-397 aa) was constructed and cloned into the PUC-sp cloning vector was procured from Sangon Biotech Shanghai, China. pET-N-GST-PreScission (Beyotime Biotechnology, Shanghai, China) was used as an expression vector for the insertion of restricted digestion gene encoding SETDB1. The recombinant vector was transferred to *E. coli* BL21 cells were cultured in LB media, the expression was induced by the addition of 0.5 mM Isopropyl beta-D-thiogalactopyranoside (IPTG) and incubated for 16 h at 16°C. The expressed protein was purified by harvesting the cells and resuspended in 0.01M phosphate-buffered saline (PBS; pH 7.4) supplemented with 2mM beta-mercaptoethanol, 5 % glycerol, 0.1 % Triton X-100 and 1 mM DL- Dithiothreitol and the cells were disrupted with incubation of Lysozyme (1 mg/mL) for 30 min. Further, the cells were lysed through ultrasonication and centrifuge at 13400 g for 30 min. The collected supernatant used as a protein source for the purification of SETDB1 by affinity chromatography using a GST-tagged resin column (Beyotime Biotechnology, Shanghai, China).

### 2.9. Circular Dichroism Spectroscopy

CD spectroscopy is one of the sensitive spectroscopy techniques to elucidate the protein secondary structure conformation upon ligand binding. The potential binding of selective compound, ECN with the target protein, SETDB1 was recorded CD spectra using JASCO J-810 CD spectropolarimeter and quartz cell with 0.1 cm and data being collected at every nanometer from 190 to 260 nm. The CD spectrum was collected with fixed protein concentration, 2 µM in tris-buffer (50 mM Tris-HCl; 150 mM NaoH; pH 7.6) and various concentrations of (5, 10 and 20 μM) compound, ECN. Three replicate experiments were carried out. The α-helixes, β-sheets, turns and random coil contents were analyzed by using Spectral manager ver.2.

## 3. Results

### 3.1. Identification of histone methylation inhibitors in the cells

Epigenetic regulation of genes through trimethylation of H3K4 and H3K9 played a vital role in biological process such as cell survival, cellular differentiation, and development. The level of histone trimethylation at H3K4 and H3K9 correlates with cell survival [36]. We found that there were seven compounds including emetine (EMT), methylcaconitin (MLA), streptonigrin (SYN), brefeldin A (BFA), harringtonine (HRT), echinomycin (ECN), dehydroandrographolide (DHA) among the 502 natural compounds in our library reduced the cell viability by more than 40% (Fig. 1A, **Table S1**). The reduced cell growth upon drug treatment were also confirmed with microscopic imaging and cell counting assay (Fig. 1B, 1C). To further investigate the inhibitory effects of these compounds on histone methylations, we performed Western blotting quantify the levels of H3K4me3 and H3K9me3 in the cells upon drug treatments. We found that the compounds ECN, SYN, and EMT showed significant inhibitions on the cellular level of H3K4me3 when compared to the control, whereas the compounds MLA, BFA, DHA, and HRT didn’t have the effects (Fig. 1D). Moreover, the compounds ECN, SYN, EMT and MLA significantly inhibited on the cellular level of H3K9me3 but BFA, DHA, and HRT did not (Fig. 1E). Our results suggest that echinomycin, emetine, and streptonigrin consistently show the biological effects on the cell growth and the H3K4 and H3K9 methylations, and may serve as candidate compounds for the downstream in silico analysis of molecular docking and molecular dynamics simulations.

**Fig. 1.**
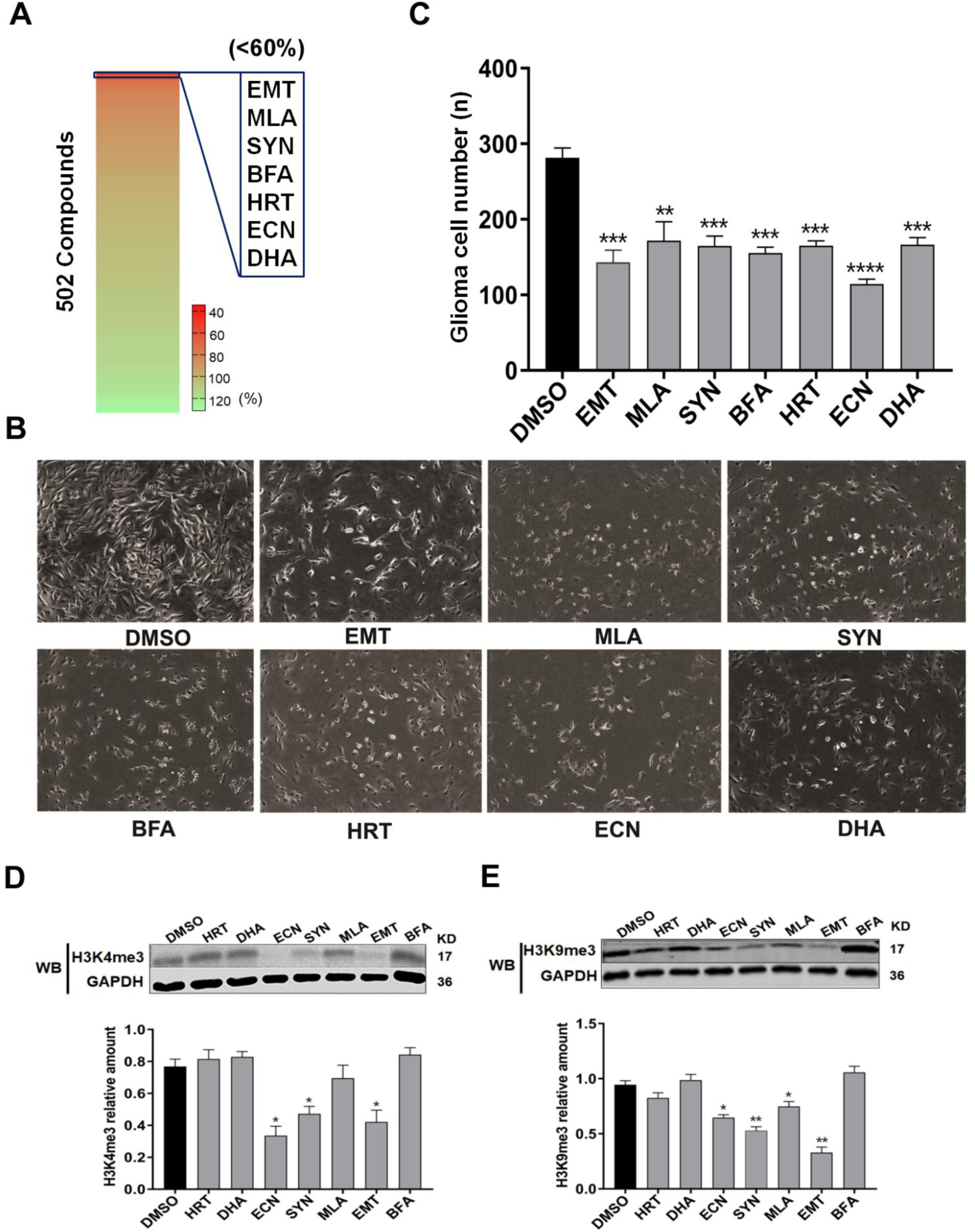
Heatmap representation of cell viability percentages of 502 natural compounds from our local dataset against U251 glioma cells, and best hits were highlighted on the top with red color (**A**), Induction of cell death by the seven compounds after 48 h incubation with the concentration of 20 μg/mL at 37 ℃ in an atmosphere of 5% CO2 (**B**). The cell count of glioma cells after 48 h incubation with these seven compounds was summarized in the bar graph (**C**). The experiment was performed three times and all the values were reported as mean±SD. The groups that have assigned **** (p<0.0001); *** (p<0.001); **(p<0.01) were statistically significant when compared to control. Expression levels of H3K4me3 (**D**), H3K9me3 (**E**) in glioma U251 cells after 48 hr incubation with seven compounds (vehicle control, DMSO) in the concentration of 20 μg/mL at 37 ℃ in an atmosphere of 5% CO2. The reference protein GAPDH was used as a loading control. The related amount of H3K4me3 and H3K9me3 was quantified by comparing with expression levels of GAPDH by Image J software and plotted as a bar graph. Error bars indicate SD from three replicates. The groups that have assigned **(p<0.01); *(p<0.05) were statistically significant when compared to control.

### 3.2. Protein-Ligand interactions by docking analysis

To elucidate the protein-ligand interactions and binding orientations, we performed molecular docking analysis of the seven compounds emetine (EMT), methylcaconitin (MLA), streptonigrin (SYN), brefeldin A (BFA), harringtonine (HRT), echinomycin (ECN) and dehydroandrographolide (DHA) with various histone lysine methyltransferases involved in H3K4 and H3K9 methylations. The crystal structures of the SET domain of all HMTases were retrieved from protein databank, which are complexes with Zn^2+^ ions, SAM and/or substrate H3 peptide. The H3 peptide binding region was chosen for docking analysis hence its pivotal role in the enzymatic catalysis. The bonding characterization including dock score [S], binding affinity [Pki], binding energy [kcal/mol] and bonding interactions of these compounds with various HMTase were computed and summarized in the Table S2, S3. The docking score values were depicted in the heatmap (Fig. 2A, B). The docking analysis showed that the compounds echinomycin, emetine and streptonigrin have better binding mode with HMTases. The 2D structures of these compounds were depicted in the Figure 2C. Furthermore, the results of docking analysis demonstrate that H3K4 HMTase NSD3 interacts with ECH and SYN while both SMYD2 and MLL3 interact with ECN. Moreover, H3K9 HMTase SETDB1, G9a and GLP have shown strong interactions with EMT and ECN. The HMTase NSD3 and SETDB1were chosen for the comprehensive studies on binding interactions and structural dynamics with the compounds due to their higher docking values.

**Fig. 2.**
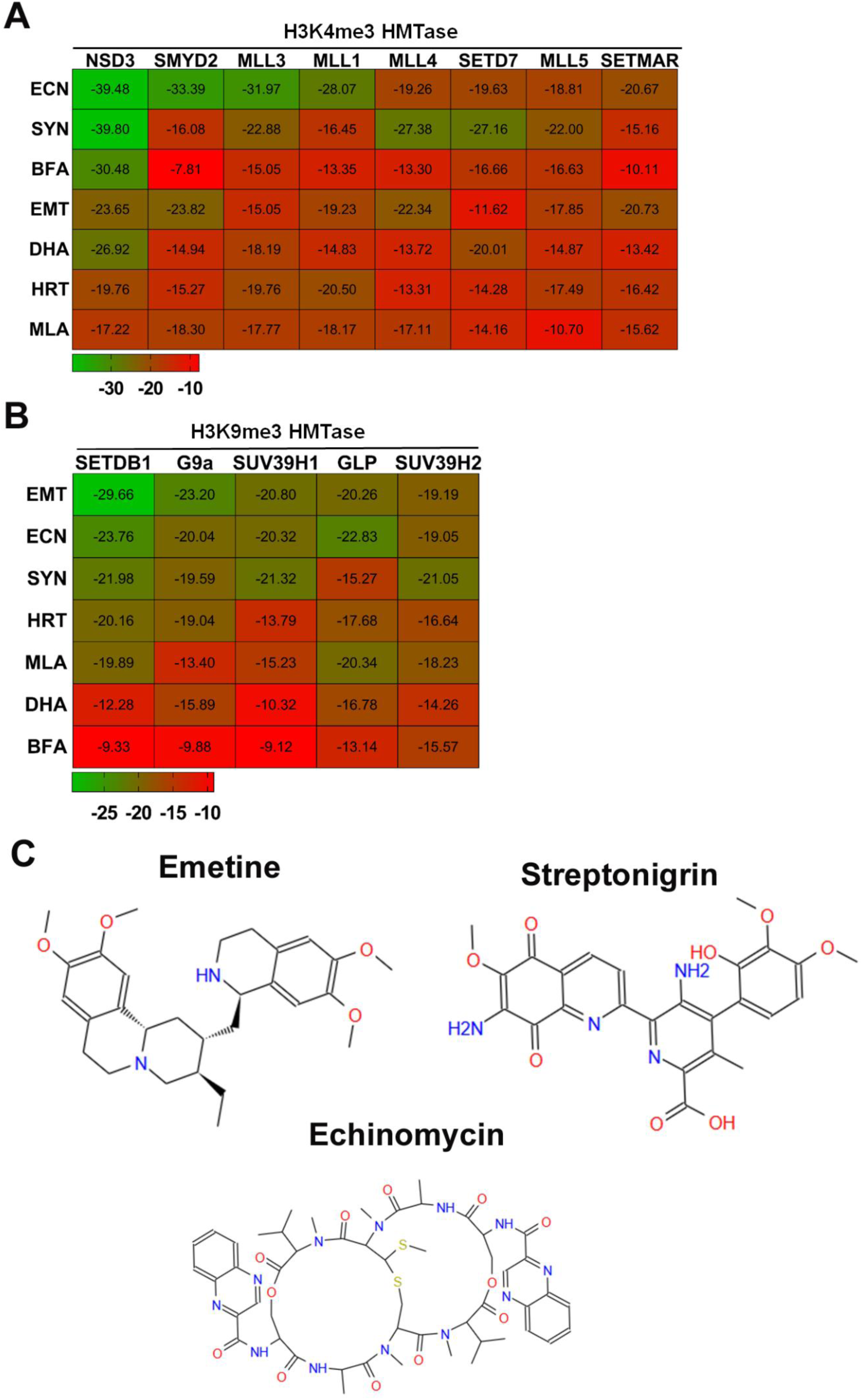
Heatmap representation of dockscore. Heatmap representation of dockscore for selective compounds against H3K4 (**A**) and H3K9 (**B**) methylation enzymes. The 2D structures of emetine, streptonigrin and echinomycin (**C**).

For NSD3, the compounds SYN and ECN have shown best dock scores of −39.80 and −39.48 respectively. The compound SYN has formed two H-bonds and one ionic bond. The carbonyl oxygen atom has formed H-bond with NE2 of His 1383 by accepting an electron pair while the carboxylic oxygen atom has formed H-bond with NE2 of His 1408 by donating an electron pair (Fig 3A) The compound ECN has formed three H-bonds and two ionic bonds. The oxygen atom of the carbonyl group acts as an H-bond acceptor forming one H-bond with NE2 of His 1383. The hydrogen atom of NH2 group and nitrogen atom within the carbon skeleton act as H-bond donor and acceptor, respectively forming H-bond with OG of Ser 1406 while the oxygen atom of the carbonyl group has formed two ionic bonds with Zn^2+^ (Fig. 3B).

**Fig. 3.**
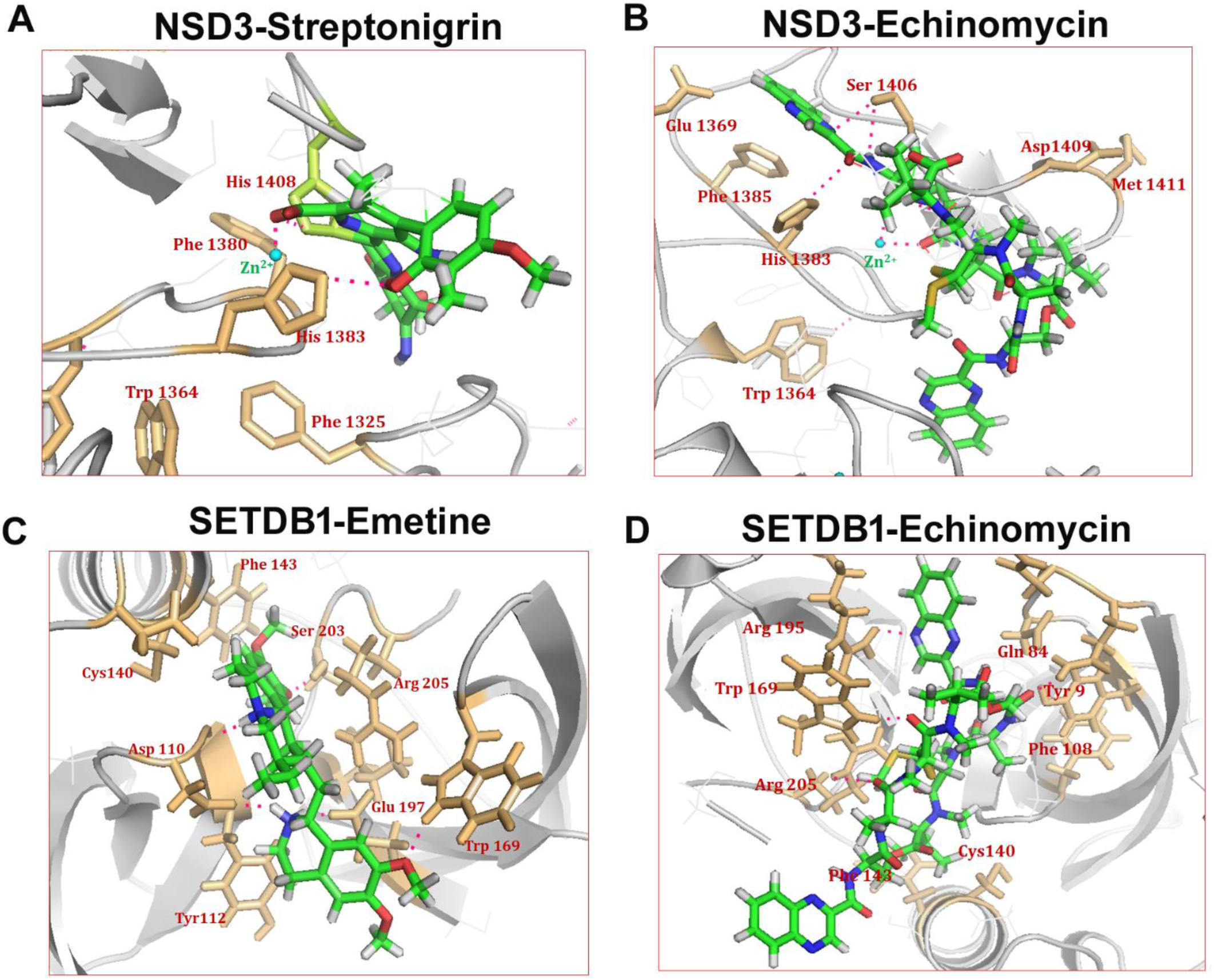
Molecular interactions of compounds with HMTases. Molecular interactions of Echinomycin and Streptonigrin with active site residues of NSD3 (**A, B**), Echinomycin with binding pocket of SMYD2 (**C**), MLL3 (**D**), GLP (**F**) and bonding interactions of Emetine with SETDB1(**E**) were captured using PyMol visualizer. All the active site residues, Zn2+ ions and H-bonds were highlighted in light orange, cyan and pink color, respectively.

For SETDB1, the EMT has interacted with the Tudor domain region and formed three H-bonds by donating an electron pair and two H-bonds by accepting an electron pair. Two amine hydrogens in EMT act as an H-bond donor forming H-bond with OD2 and the carbonyl oxygen of Asp 110. The amine hydrogen has formed H-bond with OE2 of Glu 197 by donating an electron pair. One of the carbonyl oxygens in the compound has formed H-bond with NE of Trp 169 while another one has formed H-bond with OG of Ser 203 (Fig. 3C).

The compound ECN have shown the dockscore −23.76 and formed four H-bonds by accepting an electron pair. Two oxygen atoms of EMT forming H-bond with nitrogen atom of Glu 84 and Trp 169, respectively. The nitrogen and oxygen atom of EMT forming H-bond with NH group of Arg 195 and Arg 205, respectively (Fig. 3D). These findings prompted us to carry out MD simulations of docked protein-ligand complex to elucidate the dynamic behavior of the compounds against the NSD3 and SETDB1.

### 3.3. Molecular dynamics (MD) Simulations

The structural stability and dynamic behavior were demonstrated with MD simulations of protein, protein-ligand and protein-protein complexes in the aqueous solution with a comprehensive analysis of dynamic trajectories including root mean square deviation (RMSD), RMS fluctuations (RMSF), radius of gyration (Rg) and solvent accessible surface area (SASA). The MD analysis was carried out with the HMTase NSD3 and SETDB1 since no standard reference drugs available to validate the mode of inhibition. Hence, the molecular dynamics is a pivotal approach for the detailed investigation of structural dynamics during the compound interaction.

For the structure of NSD3, the results were plotted from the calculated root mean square deviation (RMSD), RMS fluctuation (RMSF) and radius of gyration for the backbone Cα atoms of NSD3 (NSD3a), NSD3-H3K4 peptide (NSD3b), NSD3-ECN (NSD3c) and NSD3-SYN (NSD3d) complex with respect to initial model during 50 ns of MDS (Fig. 4A). From the RMSD plot, the native structure of NSD3 showed a sharp increase up to 5 ns and reached 0.3 nm by following with equilibrium in the range of 0.3-04 nm throughout the 50 ns. The NSD3b has shown a sharp increase to 0.25 nm and a further gradual increase to 0.4 nm at 15 ns followed with equilibrium throughout the 50 ns. The NSD3c and NSD3d have shown a starting increase of rmsd up to 0.4 nm at 9 ns and fluctuated within the range of 0.3 to 0.58 nm up to 30 ns then increased to 0.63 nm at 35 ns. Subsequently, the complex gets stability and equilibrated up to 50 ns (Fig. 4A).

**Fig. 4.**
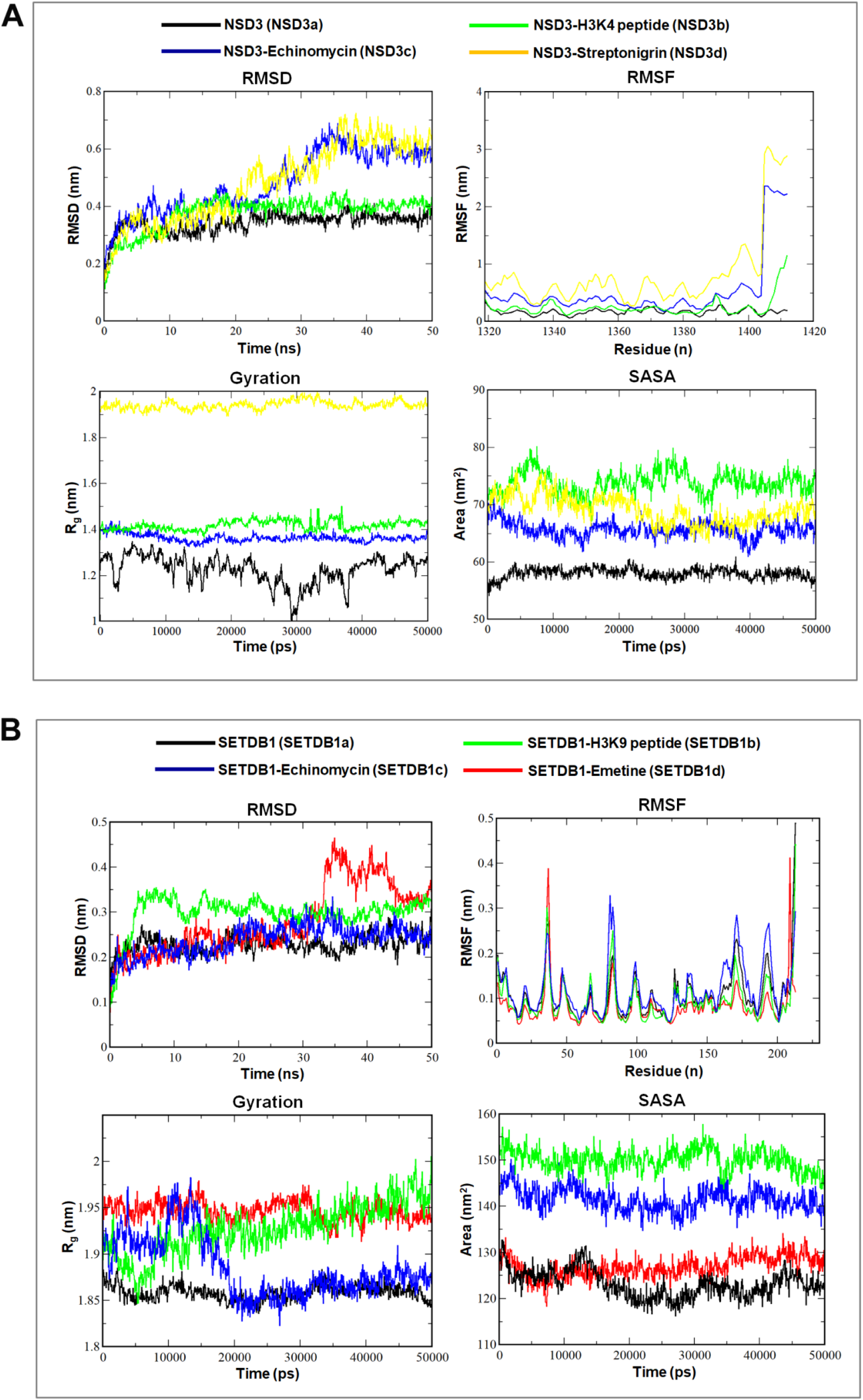
MD trajectory plots for NSD3 and SETDB1. The backbone Cα atoms root mean square deviation (RMSD) values, root mean square fluctuation of backbone Cα atoms, radius of gyration of backbone atoms and solvent accessible surface area (SASA) of NSD3, NSD3-H3K4 peptide, NSD3- Echinomycin & NSD3-Streptonigrin (**A**) and SETDB1, SETDB1-H3K9 peptide, SETDB1- Echinomycin and SETDB1-Emetine complexes (**B**) during the 50 ns time period of MD simulations.

In the same manner the dynamic trajectory analysis was performed for SETDB1 (SETDB1a), SETDB1-H3K9 peptide (SETDB1b), SETDB1-ECN (SETDB1c), and SETDB1-EMT (SETDB1d) complex with respect to initial model during 50 ns (Fig. 4B).The RMSD plot of SETDB1a has shown a sharp increase up to 0.25 nm at 5 ns followed by equilibrium with the rmsd of ~0.25 nm up to 50 ns. The SETDB1b has shown higher rmsd with a sharp increase up to 0.35 nm at 5 ns. Further, the structure has shown equilibrium throughout the 50 ns of MDS with slight fluctuations within the range of 0.28-0.35 nm. The structure SETDB1c has shown a slow increase of rmsd up to 0.25 nm at 30 ns followed by equilibrium throughout the 50 ns. While the structure SETDB1d has shown a slow increase of rmsd up to 0.28 nm at 30 ns. Further, the structure has shown sharp increase of rmsd up to 0.45 nm at 30 ns followed by gradual decrease of rmsd up to 50 ns of MDS (Fig. 4B).

The atomic RMS fluctuations were investigated for unliganded, protein-ligand and protein-protein complex. This depicted that the NSD3d and NSD3c had higher atomic fluctuations that reached up to 3.0 and 2.2 nm, respectively, in the catalytic region when compared to NSD3a and NSD3b. Even though some of the regions in NSD3a and NSD3b have shown the fluctuations ranging between 0.1 and 0.6 nm, they are not considered as significant ones because they appear outside the catalytic site (Fig. 4A). The RMSF plots of the native structures of SETDB1a and SETDB1b have shown higher atomic fluctuations in the residue region of 205-213, while the structures of SETDB1c and SETDB1d have shown higher fluctuations up to 0.4 nm in the residue regions of 30-40 and 75-90, respectively (Fig. 4B).

The Radius of gyration (Rg) is the parameter that defines the equilibrium of conformations in the total system in terms of compaction of protein structure with protein folding and unfolding [37]. In this MD analysis, Rg values were computed for protein, protein-ligand and protein-protein complexes during the 50 ns. Among these, the SYN-treated structure of NSD3d has shown higher Rg value in the range of 1.9 to 2.0 nm when compared to the structures of native NSD3a and the H3K4-NSD3b complex. While the ECN-treated structure of NSD3c has maintained the Rg variation in ~1.4 nm (Fig. 4A). It is evidenced that the compound interacts with NSD3 that alters the microenvironment of the protein structure with higher Rg value.

Rg plot of SETDB1 has demonstrated that the ECN-treated structure of SETDB1c showed an increase of Rg up to 1.97 nm at 15 ns accompanied by a gradual decrease to 1.85 nm at 20 ns and finally became equilibrated during the 50 ns period. The compound EMT-treated structure of SETDB1d has a higher Rg value with 1.95 nm and maintains the equilibrium with slight fluctuations during the 50 ns period. The structure of H3K9-SETDB1b complex has begun with Rg value 1.9 nm and slight decrease up to 10 ns followed by an increase of Rg value up to 2.0 nm during the 50 ns period. However, the native structure of SETDB1a keeps at the low level of Rg value below 1.90 nm (Fig. 4B).

SASA (Solvent accessible surface area) is a geometric measure of protein surface interactions and spread around with the outer environment of the solvent. The SASA value area (nm^2)^ is directly proportional to the rate of amino acid in the protein exposed to the solvent environment [38]. Deviations of SASA value will change the amino acid exposure to the solvent that affect the overall conformation of the protein. The results of SASA analysis showed variation among the liganded and unliganded proteins. The unliganded protein NSD3a has shown an average area of 58 nm2 whereas NSD3b and NSD3c have exposed to solvent areas with average areas of 68 and 75 nm2, respectively (Fig. 4A). SASA plot for SETDB1 has emphasized that the native structure of SETDB1a shows the SASA value within the range of 120-130 nm2. However, the structures of SETDB1b, SETDB1c, and SETDB1d have average areas of 152, 128 and 145 nm^2^, respectively, throughout the 50 ns of the time period (Fig. 4B). Perturbations of secondary structure patterns in the native NSD3, H3K4-peptide-treated NSD3, ECN-treated NSD3 and SYN-treated NSD3 in the 50 ns of MDS were analyzed using the VMD timeline plugin (Fig. 5A). The analysis report has revealed a higher range of conformational drift of secondary structure components in the ECN-treated and SYN-treated NSD3. In the structure of NSD3d, many conformational drifts have been taken place when compared to the native structure NSD3a. Among them, more significant changes with the replacement of coils and turns to Ext. Conformation was found in the region of 1320-1325 amino acid residues from 20-50 ns. The structure of NSD3c has shown significant replacement of coils to 3_10_ helices in the region of 1361-1364 amino acid residues from 0 to 28 ns. The structure NSD3b has shown significant alterations with the replacement of turns to 3_10_ helices in the region of 1385-1391 amino acid residues during 50 ns of MDS. All these secondary structural changes were summarized in Figure 5B.

**Fig. 5.**
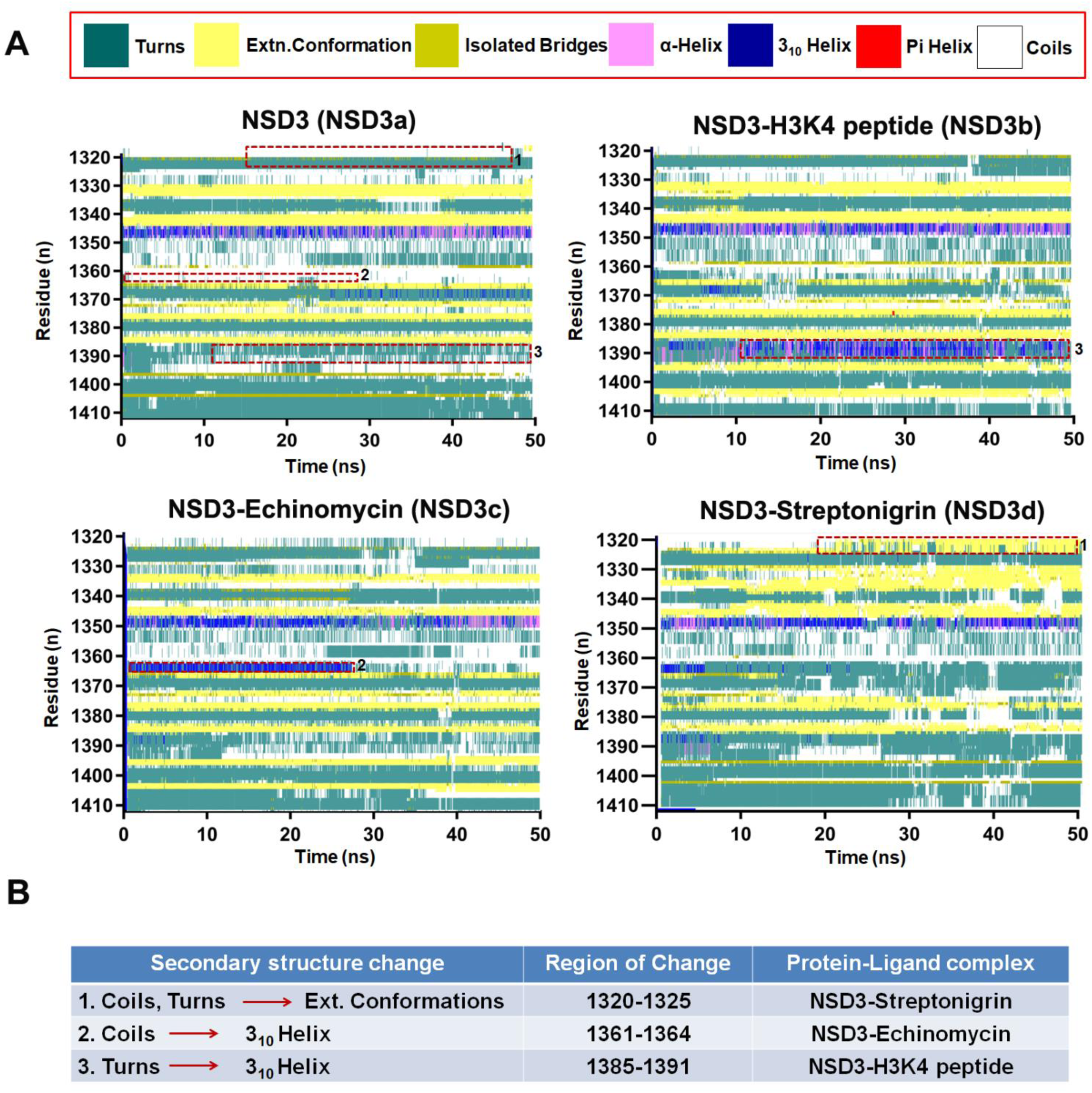
Protein secondary structure plot and summary of NSD3. Secondary structure plots of NSD3 and NSD3 interacted with H3K4, Echinomycin and Streptonigrin (**A**) during 50 ns time period of molecular dynamics simulations, reported through VMD timeline plugin. The significant sec. structural changes were highlighted with red dotted lines. The summary of secondary structure changes in the particular sequence regions (**B**).

The secondary structure plot for SETDB1 has deciphered that ECN-treated SETDB1c shows higher secondary structure alteration when compared to SETDB1a and SETDB1b. The significant changes of secondary structures were observed from 3_10_ helix to turns in the amino acid region of 74-77 and from turns to coil in the region of 205-212 (Fig. 6A). While treated with EMT, the structure of SETDB1d get a significant replacement of turns to coils in the amino acid region of 200-210 throughout the 50 ns of time period (Fig. 6A). All these secondary structural changes were summarized in Figure 6B.

**Fig. 6.**
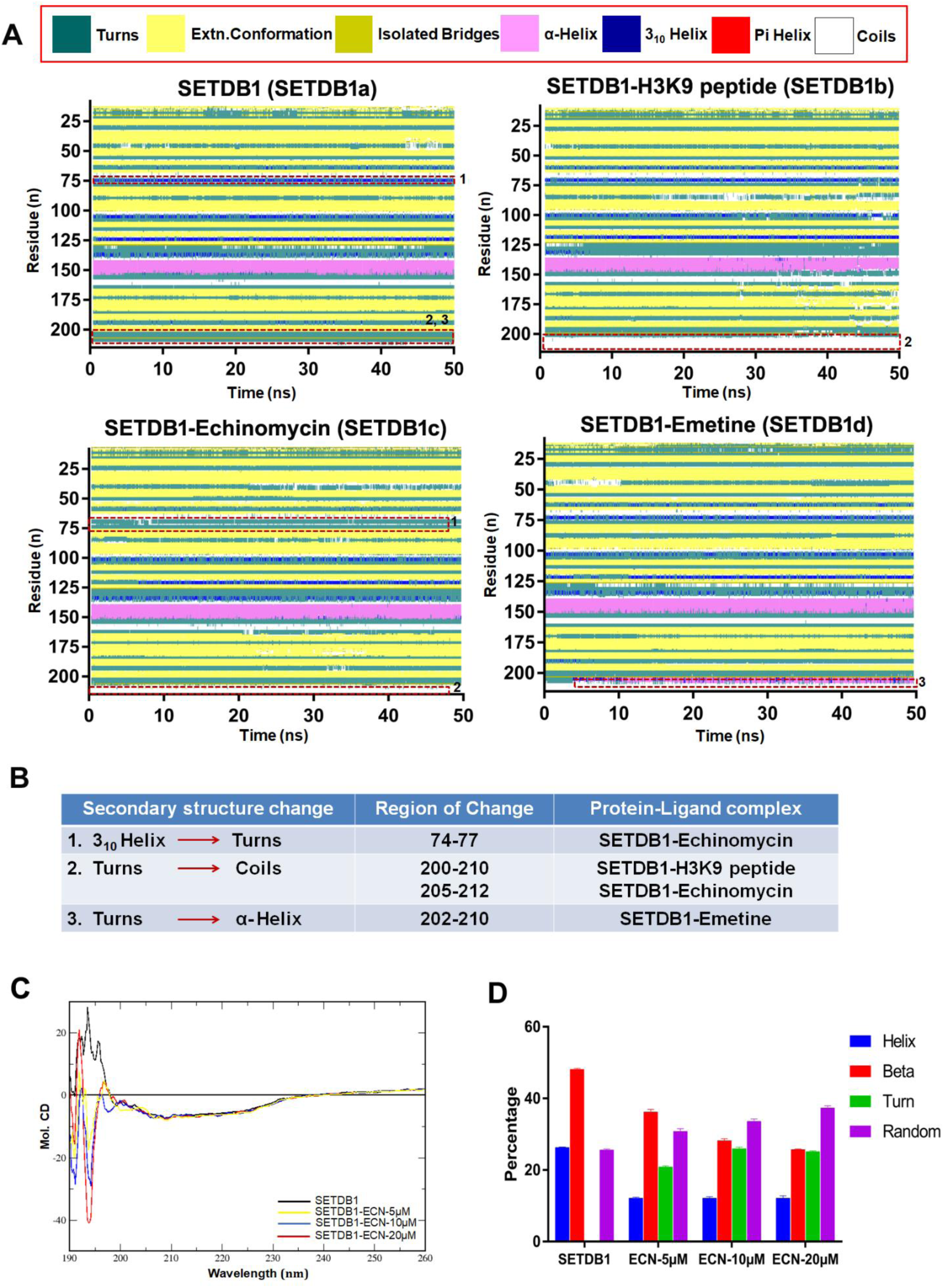
Protein secondary structure plot summary and CD spectral analysis of SETDB1. Secondary structure plots SETDB1 and SETDB1 interacted with H3K9, Echinomycin and Emetine (**A**) during 50 ns time period of molecular dynamics simulations, reported through VMD timeline plugin. The significant sec. structural changes were highlighted with red dotted lines. The summary of secondary structure changes in the particular sequence regions (**B**). Circular dichroism (CD) spectra of SETDB1 and complexes with Echinomycin (ECN) in the concentrations 5, 10 and 20 µM were recorded with JASCo-J-810 CD spectropolorimeter (C), secondary structure elements of SETDB1 and treated with ECN was calculated from its corresponding CD spectra (**D**).

Therefore, the fluctuations of number of amino acids involved in the formation of various secondary structure units in the liganded and ligand-free proteins were computed through the DSSP program. The numbers of residues and their mean±SD values at every 10 ns were plotted in the bar graphs (Fig. 7). The statistical analysis demonstrated that the turns of NSD3 increased while A-Helix decreased when interacting with ECN. However, no valid changes of NSD3 structure were observed when interacting with SYN and H3K4me0 peptide (Fig. 7A). Moreover, β-sheets and α helixes of SETDB1 decrease during the interaction with ECN. While 3-Helix of SETDB1 was found to be decreased in SETDB1 during the interaction with EMT when compared to the ligand-free protein. However, no valid changes of SETDB1 structure were observed during the interaction with H3K9me0 (Fig. 7B).

**Fig. 7.**
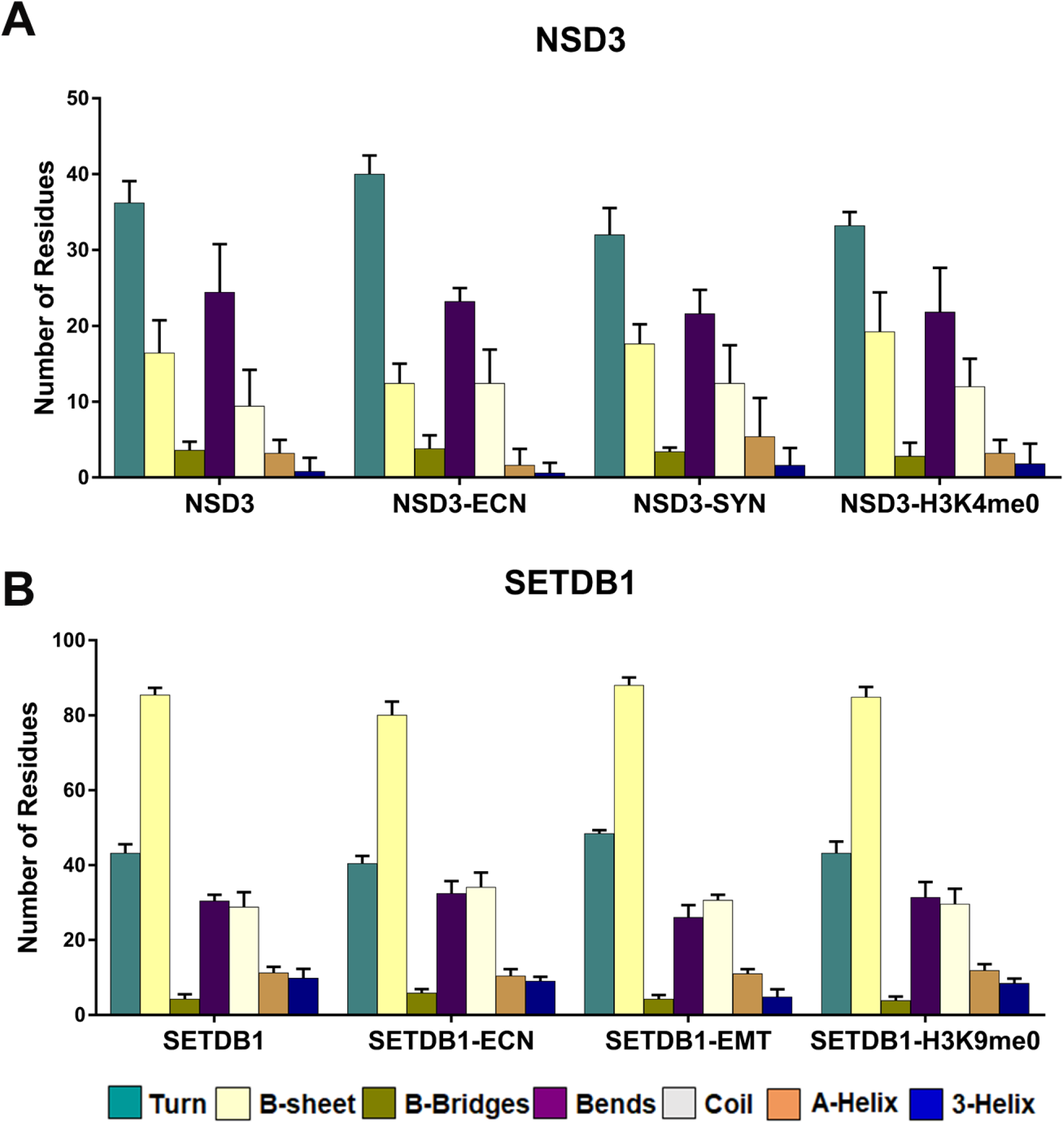
Bar graph representation of number of amino acids forming various secondary structure units. Number of amino acids involved in the formation various secondary structure units during compound and H3K4me0/H3K9me0 interacted with NSD3(**A**) and SETDB1(**B**). Error bars indicate SD from three replicates.

Apart from MD trajectory analysis, hydrogen bond stability was computed during 50 ns of simulations period for the validation of compound interactions with HMTases. The stability of hydrogen bond has been calculated between all possible donors and acceptors in the active site of protein-ligand complex with the geometric criteria of 3.4Å. The results of hydrogen bonding profile demonstrate that the maximum of six hydrogen bonds had been observed for echinomycin and streptinigrin with active site amino acids of NSD3 among these two bonds were stable and four bonds were weak. While the substrate H3K4me0 peptide has also maintained six hydrogen bonds among these three were stable (Fig. 8A). The compounds echinomycin and emetine have maintained the maximum of four and three hydrogen bonds, respectively with SETDB1. Among this one stable hydrogen bond was observed for echinomycin and emetine when compared to H3K9me0 peptide, which has maintained four hydrogen bonds among these two were stable (Fig. 8B). Together, our data suggest that echinomycin targets both NSD3 and SETDB1 while streptonigrin more specifically targets NSD3 and emetine targets SETDB1.

**Fig. 8.**
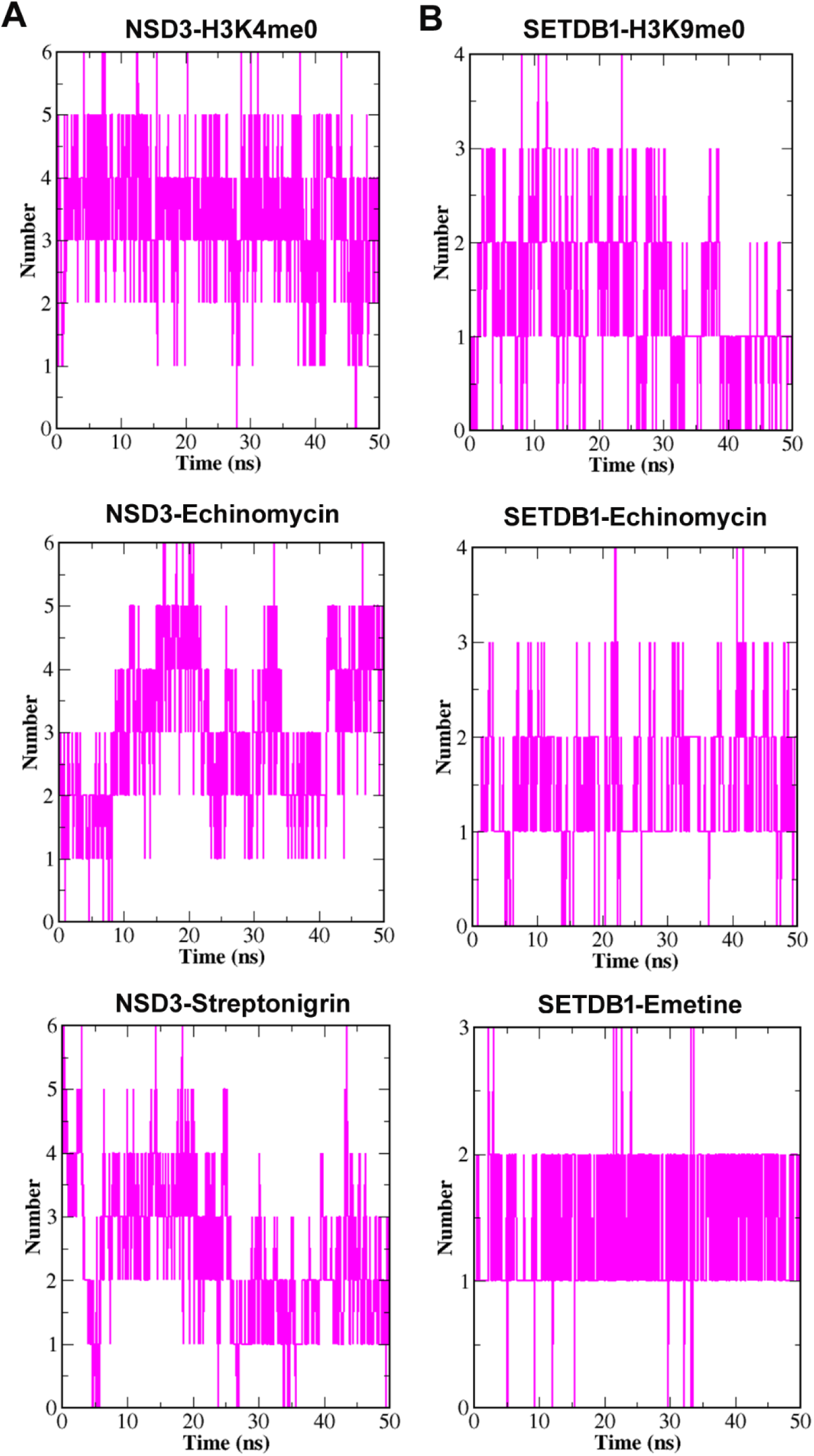
Hydrogen bond intensity of NSD3 with H3K4me0 peptide, echinomycin and streptonigrin and (A), SETDB1 H3K9me0 peptide with echinomycin and emetine (B) during 50 ns of MDS.

### 3.4. Binding free energy calculations

To get an insight in to the bioenergetics of the molecular interactions between the compounds and HMTases, the molecular mechanics-Poisson-Boltzmann Surface Area (MM/PBSA) calculations were carried out for protein-ligand complexes on the basis of last 5 ns of MD trajectory. The binding free energy of protein-ligand complex in the solvent is calculated as:

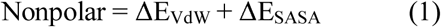

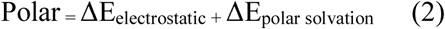

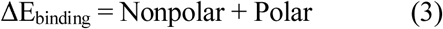

The binding energy (ΔE_binding_) was categorized in to four energy terms such as van der Waals (ΔE_VdW_), SASA (ΔE_SASA_), electrostatic (ΔE_electrostatic_) and polar solvation energies (ΔE_polar_ solvation).

In these, non-bonded interaction energies of ΔE_VdW_, ΔE_SASA_ and ΔE_electrostatic_ are pivotal and favours the stability of protein-ligand complex. Therefore, the polar solvation energy was made an opposite contribution with higher positive value then electrostatic components.

The MM/PBSA analysis results clearly demonstrated that the NSD3-Echinomycin and NSD3-Streptonigrin complexes have shown the binding free energy of −140.10 and −137.47 kJ/mol, respectively when compared to NSD3-H3K4me0 complex, which has more energy negative value of −291.58 kJ/mol (Fig. 9A). The free energy decomposition of each residue has revealed that the residues E1361, E1369 and F1380 were found in all three complexes NSD3-Echinomycin, NSD3-Streptonigrin and NSD3-H3K4me0 with the energy value of more than or equal to −20.00 kJ/mol. Preferably, the residue H1383 in the complex NSD3-Echinomycin and the residue H1408 in NSD3-Streptonigrin showed higher native value of free energy and contributes in binding interactions (Fig. 10A).

**Fig. 9.**
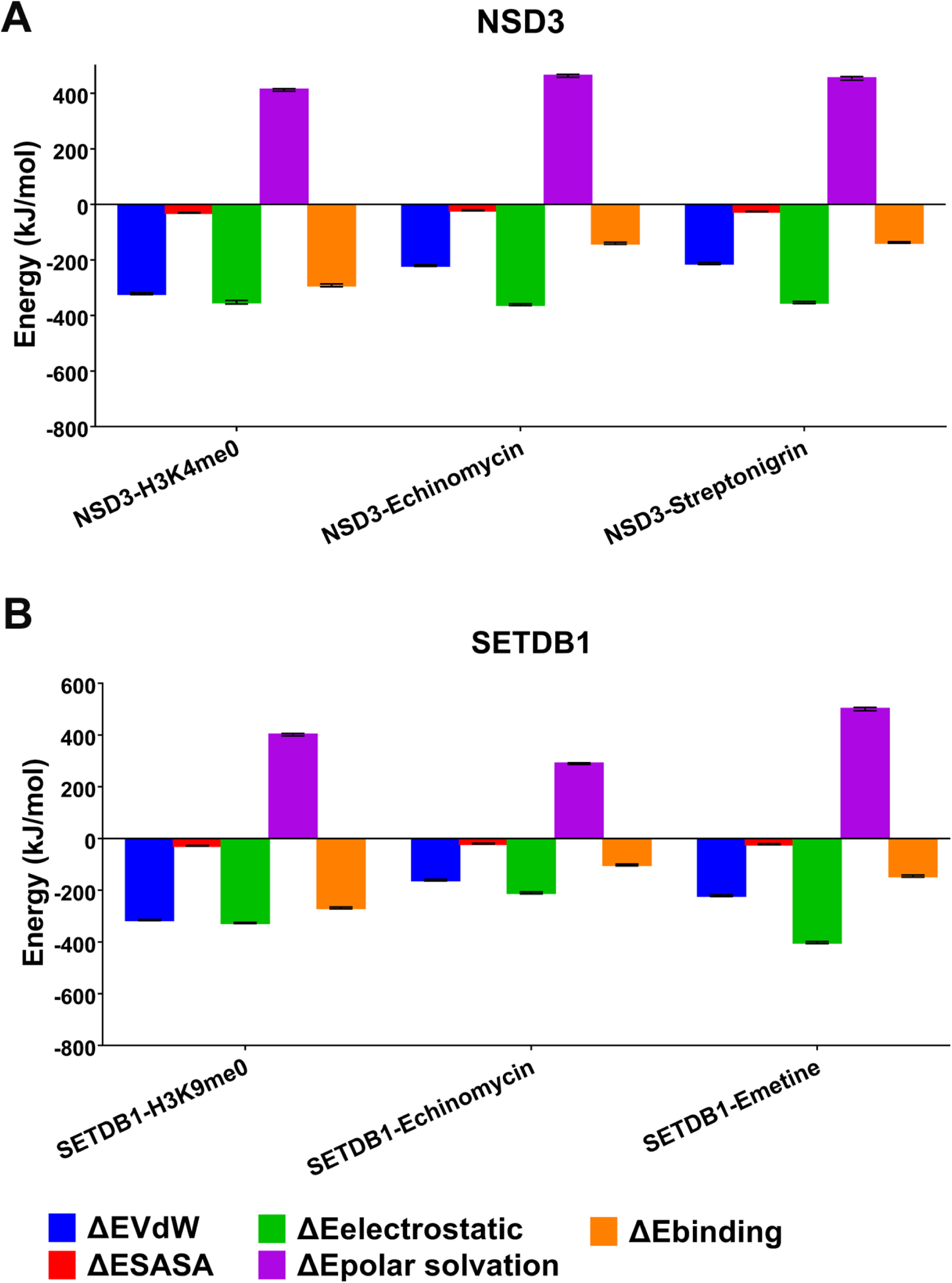
Binding free energy terms ΔEVdW, ΔESASA ΔEelectrostatic and ΔEpolar solvation of complexes NSD3- Echinomycin, NSD3-Streptonigrin and NSD3-H3K4me0 (A), SETDB2-Echinomycin, SETDB1- Emetine and SETDB1-H3K4me0 (B).

In addition, the complexes of SETDB1-Emetine and SETDB1-Echinomycin showed the binding free energy of −145.57 and −102.15 kJ/mol, respectively when compared to the complex SETDB1-H3K9me0 which has high energy native value of −268.55 kJ/mol (Fig. 9B). The residue free energy contribution plot showed that the hot residues D110, I138, W169 and S203 had binding free energy more than or equal to −20 kJ/mol and contributed for binding interactions. (Fig. 10B)

**Fig. 10.**
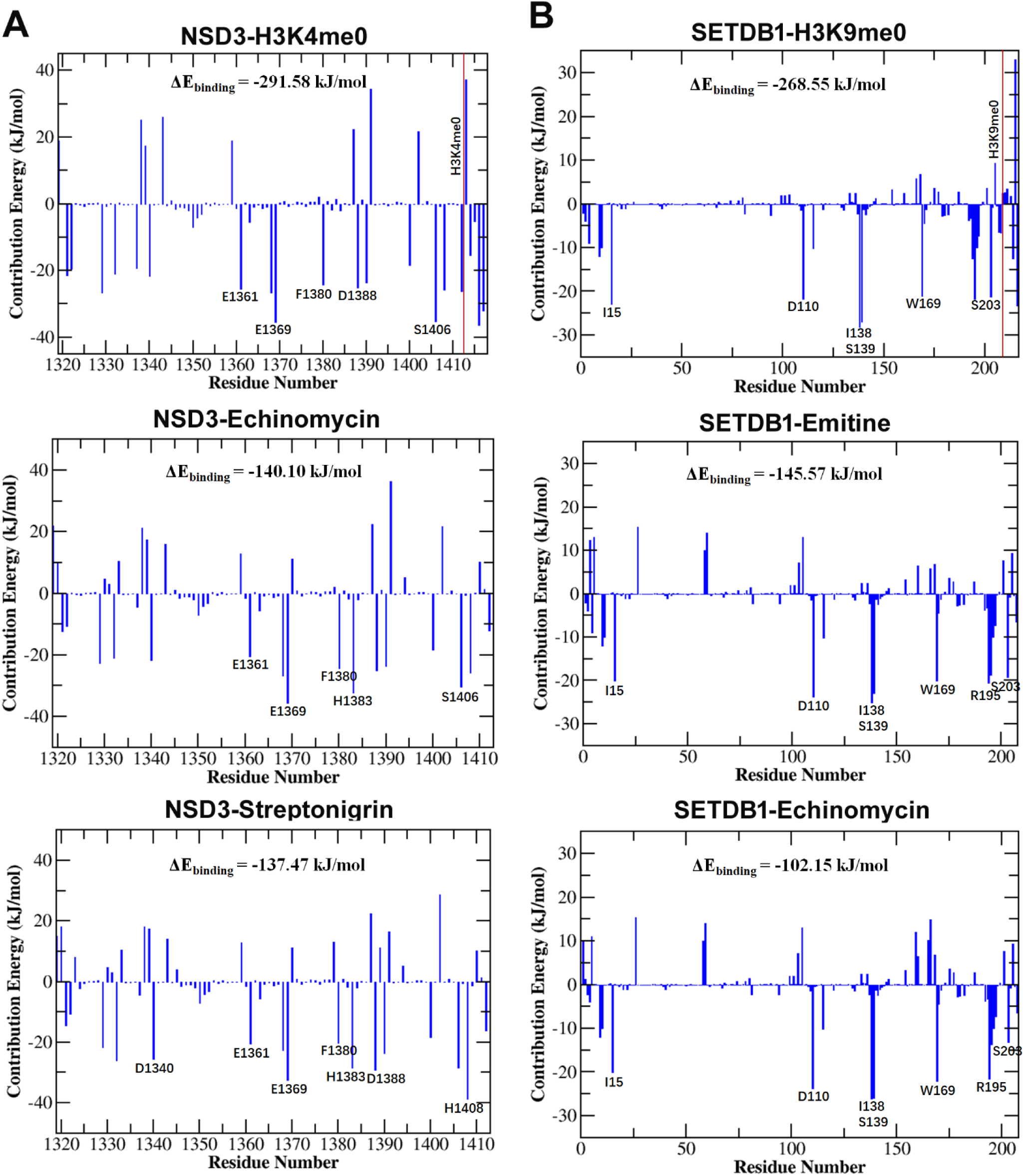
The binding free energy decomposition of each residue in the complexes NSD3-H3K4me0, NSD3-Echinomycin, NSD3-Streptonigrin (A), SETDB1-H3K4me0, SETDB2-Echinomycin, SETDB1-Emetine.

Hence, the results of binding free energy calculations and amino acid decomposition demonstrate that the NSD3 interact with ECN and SYN through the possible binding residues including E1361, E1369, F1380, H1383, D1388, S1406 and H1408. In the same way SETDB1 interact with EMT and ECN by the possible residues including D110, I138, S139, W169R195 and S203 as evidence by protein-ligand interactions from molecular docking studies.

### 3.5. Confirmation of protein-ligand interaction with Circular dichroism (CD) spectroscopy

CD spectroscopy is the comprehensive technique to analyze secondary structural alterations of the protein upon treatment with ligand molecules. In the present experiment, one of the prominent therapeutic targets SETDB1 was selected for CD analysis due to its highest binding affinity with selective ligand molecules. The SETDB1 synthetic gene was cloned and expressed in the *E. coli* system and the protein sample was purified using affinity chromatography for the assessment of CD analysis (**Fig. S1**). The compound ECN was selected due to its relatively higher inhibition on the growth of glioma cells and its significant dock score with SETDB1.

The CD spectral analysis of SETDB1 has demonstrated that the intensity of negative band at 194 nm increased in response to higher concentration of ECN (Fig. 6C). Thus, it is evidenced that ECN destabilizes the β-sheet formation when its concentration is increased. The secondary structural components in the free and compound-treated protein were calculated using the Spectral Manager version 2. The native structure of SETDB1 consists of ~26.3% α-helixes, ~48.1% β-sheets, ~25.6% random coils and 0% of turns. However, the secondary structure of compound-treated SETDB1 changes when added with 5, 10 and 20 μM of ECN, respectively. At 20 μM, the α-helixes and β-sheets were decreased to ~12.20% and ~28.2%, respectively, while the turns were increased from 0 to ~25.95% and random coils were increased from ~25.6% to ~33.65% (Fig. 6D). Hence, we revealed that the binding of ECN with SETDB1 induces the conformational changes in the secondary structure that might inactivate the catalytic property.

## 4. Discussion

Histone modifications are the significant regulators of gene expressions and disease progress. Methylated H3K4 and H3K9 are two major chromatin hallmarks to enumerate the catalytic functions of HMTases. HMTase-mediated methylations of H3K4 and H3K9 modulate the structure of chromatin and regulate the expression of downstream genes involved in the cancer [39]. These epigenetic alterations may lay the fundamental changes of transcriptions of oncogenes and/or tumor suppressor genes [40, 41]. The trimethylation of H3K4 at promoters may lead to open chromatin and gene activation, whereas trimethylation of H3K9 may cause gene silence [42, 43]. Predominantly H3K9me3 serves as a biomarker in therapeutic interventions because it is up-regulated in glioma cells and prevents apoptosis by suppressing apoptotic activators which are functionally associated with the pathogenesis of glioblastoma [44]. Keeping in view the significance of HMTase as an ideal therapeutic drug target in epigenetics, we focused on the effects of epigenetic repression by HMTases during the interactions with small molecules.

In this scenario, we used in-house library of 502 compounds for the screening of potent inhibitors of histone methylation in the cells with initial broad range of cytotoxic assay. The cytotoxicity results exemplified that the compounds EMT, MLA, SYN and BFA (> 50% cell viability) and HRT, ECN and DHA (50- 60% cell viability) were identified as potent cytotoxic compounds with possible effects on histone methylation

Hence, the scrutinized potent cytotoxic compounds emphasize for the promising screening of natural product-based leads with specific epigenetic targets. However, none of these natural compounds have been shown to target epigenetic machineries in the cells. Therefore, the seven compounds were evaluated for their effect on histone methylation in the cells. The western blot analysis revealed that the compounds ECN, EMT and SYN have shown significant inhibition of H3K4me3, while the compounds EMT, SYN, ECN, and MLA have shown inhibition of H3K9me3 levels in the cell. Based on the western blot results we hypothesized that the compounds showed effect on histone methylation cytotoxic with proven cytotoxic activity might be interact with HMTases to regulate epigenetic marks.

Furthermore, the computational techniques such as molecular docking and molecular dynamics played a pivotal role to identify new molecular entities for epigenetic targets [45]. Prospective docking screening and molecular dynamics simulations have been carried out to identify epigenetic modulating compounds and to explain mechanism of action, binding mode and protein dynamics [46, 47]. The protein-ligand interactions by docking analysis have emphasized that the HMTases NSD3, SMYD2, and MLL3 involved in H3K4 methylation highly interact with ECN and SYN. While SETDB1, GLP, G9a, and SUV39H1 in H3K9 methylation have shown strong bindings with EMT, ECN, and SYN. The MD analysis (including RMSD, RMSF, Rg, SASA and secondary structure components) of complexes of NSD3-ECN/SYN and SETDB1-EMT/ECN have shown strong bindings between the HMTases and compounds. The MD analysis also confirms that SYN preferentially binds to NSD3, while EMT preferentially binds SETDB1. ECN may act as a common H3K4 and H3K9 methylation inhibitor by interacting with both NSD3 and SETDB1.

The prediction of binding free energy of ligand with macromolecules have significant practical values in the screening of potential therapeutic drugs [48]. Molecular dynamics free energy perturbations with explicit solvent molecules through MM/PBSA method offers the most power full and promising approach to estimate free energies of protein-ligand complexes [49, 50] Our study has employed MM/PDBS calculations to enumerate the bioenergetics of NSD3 and SETDB1 with the lead compounds. The results reveal that the complex NSD3-Echinomycin and NSD3-Streptonigrin showed van der Waals (ΔE_VdW_ −220.32 and −212.32 kJ/mol), electrostatic (ΔE_elec_ −361.12 and −353.12 kJ/mol) interactions are the dominant forces that are involved in the stabilization of complexes. In addition, the complexes SETDB1-Emetine and SETDB1-Echinomycin have shown the van der Waals (ΔE_VdW_ −221.38 and −161.12 kJ/mol), electrostatic (ΔE_elec_ −402.30 and −210.33 kJ/mol). Therefore, the residue decomposition analysis suggested that the residues H1383 and S1405 showed more free energy values −32.41 and −30.3 kJ/mol, respectively in the NSD3-Echinomycin complex and formed stable hydrogen bond interactions as shown in the Figure 3B. In the NSD3-Streptonigrin complex the residues H1383 and H1408 contribute in the hydrogen bond interactions with the free energies of −24.40 and −25.74 kJ/mol, respectively (Fig. 10A) as shown in the Figure 3A. In addition, the residue decomposition of SETDB1 has demonstrated that the residues D110, I138, W169 and R195 in the complexes SETDB1-Emetine and SETDB1-Echinomycin showed binding free energy more than or equal to −20 kJ/mol. Among these the residues D110, W169 and R195 have formed stable hydrogen bond interactions as depicted in the Figure 3C, D

Hence, the ensembled molecular docking and molecular dynamics trajectories followed by MM/PBSA calculations revealed that these natural compounds bound to HMTases and potentially inhibited the catalytic functions of HMTases. Collectively, our study revealed that echinomycin, emetine, and streptonigrin exhibit pivotal interactions with the HMTases. Our investigation provides a novel platform and a starting point to develop a new class of natural-product based drugs as persuaded potent HMTase inhibitors which can be considered as a promising lead in the epigenetic drug discovery.

## Acknowledgments

We are grateful to the members of the Huang lab, especially Meiyang Li, Cuicui Yu, Bo Wang, and Hongzhi Guo for technical assistance and helpful comments.

