## Supplemental Figures and Tables for "Discovery of potential epigenetic inhibitors against histone methyltransferases through molecular docking and molecular dynamics simulations"

**Supplementary Material**

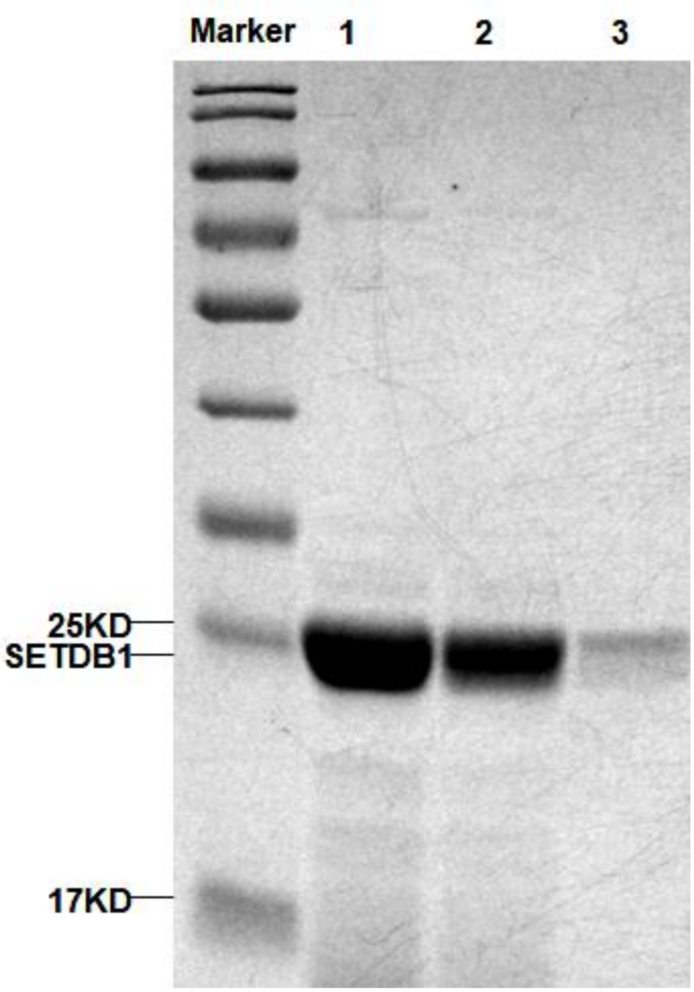


**Figure S1.** Coomassie bright blue staining of purified protein, SETDB1 (23 kD), Lane 1, 2 and 3 are liquid flow outs 1, 2 and 3 times from the column

**Table S1.** Cell viability percentages of 502 natural compounds against U251 glioma cells

| **Compound Name** | **Cell viability (%)** | **Compound Name** | **Cell viability (%)** |
|---|---|---|---|
| **Emetine** | **34.46** | Bicuculline, (+)- | 75.80 |
| **Methyllycaconitine citrate** | **44.15** | Butein | 76.09 |
| **Streptonigrin** | **49.91** | Rauwolscine | 76.29 |
| **Brefeldin A** | **49.94** | Deltaline | 76.93 |
| **Harringtonine** | **50.79** | Eburnamonine, (-)- | 77.05 |
| **Echinomycin** | **57.97** | Salsolinol HBr | 77.21 |
| **Dehydroandrographolide** | **58.25** | Pseudopelletierin HCl | 78.16 |
| Nonactin | 60.74 | Quinidine HCl | 78.87 |
| Vinblastine sulfate | 63.91 | Rotenone | 79.18 |
| Antimycin A1 | 66.14 | Delcorine | 79.19 |
| Thapsigargin | 66.72 | Brucine n-oxide | 79.31 |
| Taxol | 67.43 | Strychnine HCl | 79.34 |
| Radicicol | 67.80 | Rottlerin | 79.98 |
| Vincristine sulfate | 69.39 | Resveratrol | 80.08 |
| Phorbol 12-myristate 13-acetate, 4-a - | 73.81 | Eriodictyol | 80.38 |
| Tunicamycin B | 74.38 | Sterigmatocystin | 81.05 |
| Sitosterol, b - | 74.46 | Arecoline HBr | 81.16 |
| Kaempferol | 74.61 | Emodin | 81.39 |
| Anisomycin | 75.32 | Gramine | 81.52 |
| Rosmarinic acid | 75.79 | Veratridine | 81.59 |
| Veratramine | 81.71 | Condelphine | 84.56 |
| Eriocitrin | 81.71 | Robinetine | 84.58 |
| Narasin | 82.20 | Quassin | 84.80 |
| Bavachinin A | 82.27 | Quercetin | 84.86 |
| Actinomycin D | 82.39 | Diacetylkorseveriline | 84.94 |
| Harmaline HCl | 82.48 | Cotinine, (-)- | 85.01 |
| Chrysoeriol | 82.67 | Chaetomellic acid A | 85.07 |
| Lysergol | 82.69 | Rhamnetine | 85.14 |
| Desoxypeganine HCl | 83.13 | Austricin | 85.48 |
| Datiscetin | 83.15 | Phytosphingosine | 85.74 |
| Harmine HCl | 83.42 | Rifampicin | 85.76 |

| **Compound Name** | **Cell viability (%)** | **Compound Name** | **Cell viability (%)** |
|---|---|---|---|
| Catalpol | 83.44 | Hyoscyamine | 85.82 |
| Convolvamine HCl | 83.56 | Pilocarpine HCl | 85.85 |
| Apigenin-7-O-glucoside | 83.85 | Deacetylcolchicine, N-formyl- | 85.89 |
| Kanamycin | 83.98 | Kahweol | 86.08 |
| Chrysine | 83.98 | Wortmannin | 86.13 |
| Prostaglandin J2 | 84.01 | Eudesmine | 86.15 |
| Homobutein | 84.37 | Podophylotoxin | 86.37 |
| Oxytetracycline, a -apo- | 84.54 | Dihydroxy-4'-methoxy-chalcone, 2',6'- | 87.05 |
| Dihydroergotamine methanesulfonate | 84.55 | Vinpocetine | 87.15 |
| Berbamine 2HCl | 87.21 | Atropine sulfate | 89.81 |
| Apigenin | 87.25 | Solanidine | 89.86 |
| Tamarixetine | 87.46 | Ketopinic acid | 90.14 |
| Bufalin | 87.79 | Trimethylpsoralen, 4,5',8- | 90.27 |
| Wedelolactone | 87.94 | Prostaglandin E1 | 90.41 |
| Ungerine nitrate | 88.03 | Prostaglandin E2 | 90.50 |
| Mycophenolic acid | 88.23 | Monensin Na | 90.56 |
| Syrosingopine | 88.26 | Didymin | 90.57 |
| Swainsonine | 88.81 | Taxifolin (+) | 90.76 |
| Tschimganine | 88.90 | Quinine HCl | 90.81 |
| Lycorine HCl | 88.94 | Gliotoxin | 90.96 |
| Scopolomine N-butylbromide | 88.97 | Deoxyphorbol 13-phenylacetate 20-acetate, 12- | 90.97 |
| Monocrotaline | 89.07 | Retinoic acid, 13-cis- | 90.97 |
| Narirutin | 89.35 | Menadione | 91.13 |
| Daidzin | 89.38 | Daunorubicin HCl | 91.21 |
| Indirubin | 89.51 | Feroline | 91.29 |
| Tetrahydroalstonine | 89.64 | Phorbol 12-myristate 13-acetate | 91.39 |
| Deoxyphorbol 13-acetate, 12- | 89.68 | Ivermectin | 91.50 |
| Colchicine | 89.69 | Sclerotiorin | 91.64 |
| Bleomycin | 89.71 | Mithramycin A | 91.70 |
| Cytochalasin D | 91.80 | Penicillamine, L- | 94.04 |
| Rhoifolin | 91.89 | Picrotin | 94.06 |
| Camptothecin | 91.94 | Spectinomycin sulfate | 94.45 |
| Amentoflavone | 91.99 | Remerine HCl | 94.51 |
| Dubinidine | 92.02 | Genistin | 94.57 |

| Compound Name | Cell viability (%) | Compound Name | Cell viability (%) |
|---|---|---|---|
| Salinomycin | 92.11 | Protopine HCl | 94.61 |
| Baccatin III | 92.21 | Cytochalasin E | 94.69 |
| Isosakuranetin | 92.53 | Ouabain (-)- | 94.79 |
| Prostaglandin F2a | 92.73 | Demethylepipodophyllotoxin, 4'- | 94.89 |
| Vitexin-2''-O-rhamnoside | 92.84 | Aphidicolin | 94.90 |
| Chelidonine, (+)- | 92.87 | Epibatidine, (±)- | 94.98 |
| Mimosine, L- | 92.96 | Scoulerine | 95.21 |
| Rapamycin | 92.97 | Tanshinone IIA | 95.34 |
| Retinoic acid (all trans) | 93.21 | Tubercidin | 95.88 |
| Fisetin | 93.39 | Panaxadiol | 96.31 |
| Andrographolide | 93.46 | Prostaglandin A1 | 96.64 |
| Laudanosoline HBr | 93.51 | Arbutin | 96.73 |
| Kaempferol-7-neohesperidoside | 93.82 | Retinoic acid, 9-cis- | 96.76 |
| Lapachone, b - | 93.90 | Luteolin-3',7-di-o-glucoside | 96.81 |
| Dihydrotanshinone | 93.96 | Hypocrellin B | 96.88 |
| Isorhamnetine-3-O-rutinoside | 96.91 | Flavokawain B | 98.40 |
| Carminic acid | 96.94 | Fumagillin | 98.49 |
| Forskolin | 96.94 | Pratol | 98.66 |
| Luteolin | 96.98 | Dihydroxyflavone, 6,7- | 98.74 |
| Cinobufagin | 97.03 | Chromomycin A3 | 98.81 |
| Hydrocotarnine HBr | 97.13 | Solanine, a - | 98.90 |
| Isoscopoletine | 97.18 | Vitexin | 98.91 |
| Karakoline | 97.22 | Chartreusin | 99.09 |
| Theobromine | 97.28 | Etoposide | 99.33 |
| Sevedindione | 97.31 | Anisodamine | 99.36 |
| Sclareol | 97.43 | Ebelactone B | 99.73 |
| Myricitrin | 97.51 | Coumermycin A1 | 99.80 |
| Prostaglandin B1 | 97.59 | Daidzein | 99.93 |
| Ajmaline | 97.60 | Melatonin | 99.96 |
| Amygdalin | 97.75 | Manool | 99.99 |
| Kaempferol-3-O-glucoside | 97.96 | Picrotoxinin | 100.08 |
| Lincomycin | 98.23 | Geldanamycin | 100.13 |
| Maritimein | 98.23 | Isotetrandrine | 100.16 |
| Embelin | 98.25 | Dihydroergocristine mesylate | 100.17 |
| Morin | 98.31 | Nigericin Na | 100.18 |

| Compound Name | Cell viability (%) | Compound Name | Cell viability (%) |
|---|---:|---|---:|
| Kavain (+/-) | 100.20 | Vanillylacetone | 101.42 |
| Bromolaudanosine, (±)-6'- | 100.24 | Isovitexin | 101.55 |
| Tetrandrine | 100.31 | Decoyinine | 101.63 |
| Troleandomycin | 100.34 | Gelsemine HCl | 101.64 |
| Ellipticine | 100.47 | Songorine | 101.94 |
| Naringenin | 100.55 | Cevadine | 101.96 |
| Kainic acid | 100.58 | Quisqualic acid | 101.98 |
| Neomycin sulfate | 100.58 | Salsolidine | 101.99 |
| Imperialine | 100.62 | Myricetin | 102.00 |
| Jasmonic acid | 100.63 | Genistein | 102.01 |
| Lapiferine | 100.67 | Piceatannol | 102.03 |
| Ursolic acid | 100.69 | Cyclopiazonic acid | 102.10 |
| Caffeic Acid | 100.80 | Santonin | 102.10 |
| Harmol HCl | 100.87 | Shikonin | 102.14 |
| Trigonelline HCl | 101.03 | Usnic acid, (+)- | 102.17 |
| Myriocin | 101.09 | Vulpinic acid | 102.17 |
| Hydroxyflavone, 7- | 101.32 | Gingerol | 102.18 |
| Madecassic acid | 101.32 | Nitrarine 2HCl | 102.27 |
| Gibberellic acid, (+)- | 101.36 | Boldine | 102.40 |
| Betulinic acid | 101.41 | Palmatine Cl | 102.47 |
| Hydrastine, D-b - | 102.65 | Ellagic acid | 103.81 |
| Geranylgeranoic acid | 102.68 | Guaiol | 103.81 |
| Lapidine | 102.68 | Norharmane | 103.82 |
| Saponarin | 102.72 | Phloretin | 103.86 |
| Mitomycin C | 102.95 | Cycloheximide | 103.93 |
| Peruvoside | 103.06 | Marein | 103.99 |
| Bilobalide | 103.13 | Capreomycin sulfate | 103.99 |
| Osthole | 103.13 | Fumonisin B2 | 103.99 |
| Anabasine HCl | 103.18 | Carnitine Cl, (±)- | 104.07 |
| Ecdysone | 103.21 | Ginkgolide B | 104.11 |
| Heteratisine | 103.24 | Ingenol 3,20-dibenzoate | 104.13 |
| Homoorientin | 103.28 | Curcumin | 104.25 |
| Khellin | 103.30 | Mezerein | 104.26 |
| Nicotine, (-)- | 103.47 | C6 Ceramide | 104.30 |
| Methoxyflavone, 5- | 103.51 | Noreleagnine | 104.30 |

| Compound Name | Cell viability (%) | Compound Name | Cell viability (%) |
|---|---|---|---|
| Lupinine | 103.51 | Pinocembrin | 104.33 |
| Rhapontin | 103.60 | Vinorelbine base | 104.39 |
| Phorbol | 103.68 | Isorhamnetine-3-O-glucoside | 104.43 |
| Ferulic acid | 103.69 | Norfluorocurarine | 104.43 |
| Parthenolide | 103.76 | Scopolamine N-oxide HBr, (-)- | 104.48 |
| Cryptotanshinone | 104.53 | Plumbagin | 105.20 |
| C2 Phytoceramide | 104.56 | Lobeline HCl | 105.34 |
| Graveoline | 104.59 | Doxorubicin HCl | 105.39 |
| Valinomycin | 104.64 | Antibiotic A-23187 | 105.48 |
| Tschimganidin | 104.66 | Cycloserine, L- | 105.54 |
| Gambogic acid | 104.70 | Acacetine | 105.74 |
| Chlorogenic acid | 104.72 | Isorhoifoline | 105.75 |
| Himbacine | 104.74 | Corydaline | 105.83 |
| Oleanolic acid | 104.74 | Perillic acid | 106.08 |
| Epicatechin, (-)- | 104.79 | Aucubin | 106.21 |
| Harmane HCl | 104.79 | Geniposide | 106.45 |
| Hydroxycamptothecin, 10- | 104.81 | Catharanthine base | 106.50 |
| Harmalol HCl | 104.86 | Cerulenin | 106.57 |
| Puromycin | 104.86 | Salsoline | 106.69 |
| Aristolochic acid A | 104.87 | Dihydromethysticin | 106.73 |
| Gossypol | 104.95 | Rubescensin A | 106.76 |
| Trichodesmine | 104.96 | Cyclopamine | 106.81 |
| Jervine | 105.06 | Castanospermine | 106.82 |
| Methylergonovine | 105.15 | Grayanotoxin III | 106.84 |
| Bromocriptine mesylate | 105.17 | Muscarine Cl, (+)- | 106.94 |
| Amphotericin B | 106.98 | Sclareolide, (3aR)-(+)- | 108.63 |
| Cantharidin | 107.09 | Oligomycin A | 108.64 |
| Senecionine | 107.25 | Schisandrin A, R(+)- | 108.66 |
| Bulleyaconitine A | 107.27 | Mevastatin | 108.67 |
| Vineomycin A1 | 107.31 | Tetrahydrolipstatin | 108.74 |
| Epigallocatechin gallate | 107.36 | Diosmetine | 108.74 |
| Homoeriodictyol (-) | 107.44 | Domoic acid | 108.75 |
| Dicoumarol | 107.45 | Visnagin | 109.02 |
| Gitoxigenin | 107.84 | Auraptene | 109.02 |
| Cytochalasin B | 107.94 | Capsaicin | 109.10 |

| Compound Name | Cell viability (%) | Compound Name | Cell viability (%) |
|---|---:|---|---:|
| Coumestrol | 108.01 | Papaverine HCl | 109.17 |
| Indole-3-butyric acid | 108.12 | Peganole | 109.19 |
| Cyclosporin A | 108.20 | Cinchonine, (+)- | 109.23 |
| E-64-D | 108.25 | Indole-3-carbinol | 109.25 |
| Heliotrine | 108.39 | Lagochiline | 109.25 |
| Tomatidine | 108.41 | Hernandezine | 109.34 |
| Retrorsine | 108.50 | Zearalanol, b - | 109.66 |
| Deoxyshikonin | 108.53 | Baicalein | 109.66 |
| Isoquercitrine | 108.54 | Citrinin | 109.82 |
| Seneciphylline | 108.61 | Ryanodine | 109.87 |
| Phorbol 12,13-dibutyrate | 109.88 | Honokiol | 111.50 |
| Shikimic Acid | 110.01 | Scopoletin | 111.50 |
| Methysticin | 110.06 | Xanthotoxin | 111.55 |
| E-64 | 110.15 | Berberine HCl | 111.60 |
| Euphorbiasteroid | 110.31 | Oxocafestol, 16- | 111.78 |
| Physostigmine | 110.35 | Nomilin | 111.85 |
| E-64-C | 110.36 | Artemisinin | 111.87 |
| Peganine | 110.42 | CAPE | 111.97 |
| Cinchonidine, (-)- | 110.43 | Sanguinarine | 112.17 |
| Galanthamine HBr | 110.43 | Acivicin | 112.17 |
| Stachydrine HCl | 110.46 | Distamycin A | 112.23 |
| Isocorydine HCl | 110.47 | Ingenol | 112.31 |
| Demissidine | 110.49 | Kinetin | 112.36 |
| Deguelin | 110.67 | Geraldol | 112.43 |
| Huperzine A, (-)- | 110.70 | Asiatic acid | 112.62 |
| Homatropine HBr | 110.71 | Rutaecarpine | 112.69 |
| Cephradine | 110.75 | Skimmianine | 112.94 |
| Kenpaulone | 110.85 | Dipterocarpol | 113.12 |
| Chelerythrine Cl | 110.86 | Nalidixic acid | 113.14 |
| Lappaconitine | 111.02 | Diindolylmethane | 113.18 |
| Rutin | 113.19 | Noscapine, (±)- | 114.40 |
| Caryophyllene oxide | 113.20 | Pellitorine | 114.52 |
| Betulin | 113.22 | Chaconine, a - | 114.53 |
| Australine HCl | 113.29 | Fillalbin | 114.58 |
| Naringin | 113.53 | Carnosic acid | 114.61 |

| Compound Name | Cell viability (%) | Compound Name | Cell viability (%) |
|---|---:|---|---:|
| Tetrahydropapaverine | 113.65 | Artesunate | 114.64 |
| Sinensetine | 113.69 | Uvaol | 114.65 |
| Ochratoxin A | 113.73 | Isoreserpine, (-)- | 115.15 |
| Ecdysone, b- | 113.78 | Cafestol acetate | 115.22 |
| Phlorizine | 113.86 | Hypericin | 115.28 |
| Glycyrrhetinic acid, 18-b - | 113.87 | Conessine | 115.38 |
| Sinomenine | 113.89 | Cepharanthine | 115.44 |
| Hesperidine | 113.91 | Vincamine | 115.59 |
| Yohimbine HCl | 114.11 | Bergenin | 115.59 |
| Formononetin | 114.12 | Minocycline | 115.64 |
| Cotininecarboxylic acid, trans-4- | 114.14 | Hirsutine | 115.68 |
| Hypocrellin A | 114.15 | Yangonin | 115.92 |
| Ginkgolide A | 114.20 | Hesperetine | 116.13 |
| Panaxatriol | 114.22 | Vindoline | 116.18 |
| Thymoquinone | 114.33 | Azomycin | 116.31 |
| Corynanthine | 116.35 | Sedanolide | 118.68 |
| Oxokahweol, 16- | 116.62 | Piperine | 118.68 |
| Protoveratrine B | 116.75 | Solasodine | 118.93 |
| Sulfuretine | 116.80 | Diosmin | 119.33 |
| Podocarpic acid | 116.93 | Quercitrin | 119.34 |
| Magnolol | 116.97 | Lasalocid A Na | 119.36 |
| Dihydrolysergol, 9,10- | 117.11 | Griseofulvin, (+)- | 119.56 |
| Aconitine | 117.11 | Dihydrorobinetin | 119.67 |
| Cafestol | 117.13 | Sarsasapogenin | 119.74 |
| Bergapten | 117.38 | Pimaricin | 119.82 |
| Myristicin | 117.39 | Aloe-emodine | 120.46 |
| Silybine | 117.45 | Flavanomarein | 120.48 |
| Tropine | 117.45 | Ferutinin | 121.21 |
| E6 Berbamine | 117.57 | Biochanin A | 121.28 |
| Friedelin | 117.62 | Brassinin | 121.67 |
| Matrine | 117.73 | Hordenine sulfate | 122.06 |
| Solanesol | 118.27 | Reserpine | 122.06 |
| Hydroxytropinone, 6- | 118.29 | Rifamycin SV-NA | 122.83 |
| Neohesperidin | 118.38 | Wogonin | 122.90 |
| Zearalenone | 118.54 | Evodiamine | 123.08 |

| Compound Name | Cell viability (%) | Compound Name | Cell viability (%) |
|---|---:|---|---:|
| Securinine | 123.13 | Leucomisine | 125.43 |
| Strophanthidin acetate | 123.26 | Limonin | 125.64 |
| Lavendustin B | 124.04 | Schisandrin B, S(-)- | 126.21 |
| Schisantherin A | 124.34 | Asarinin, (-)- | 126.23 |
| Tryptanthrin | 124.53 | Silymarin | 126.24 |
| Syringetine-3-O-glucoside | 124.55 | D-Tubocurarine-chloride | 127.62 |
| Catechin hydrate, (+)- | 124.61 | Flavokawain A | 127.70 |
| Spartein sulfate (-)- | 124.75 | Galangine | 128.57 |
| Abscisic acid, (±)- | 124.90 | Lavendustin A | 128.86 |
| Cytisine, (-)- | 125.11 | Oxyacanthine sulfate | 130.79 |
| Dehydrokawain, 5,6- | 125.13 | Patulin | 130.97 |

**Table S2.** Bonding characterization, dock scores and binding energies of selective seven compounds against various therapeutic targets, H3K4 methyltransferase enzymes

| S.No | Compound | Dock Score (S) | Binding affinity (pki) | Binding energy (kcal/mol) MM-GB/VI | Bonding interactions | Bond length (Å) | Bond type |
|---|---|---|---|---|---|---|---|
| | **Histone-lysine N-methyltransferase SETMAR** | | | | | | |
| 1 | Echinomycin | -20.67 | 7.67 | -32.26 | O------N Val 191 | 2.59 | H-acc |
| 2 | Streptonigrin | -15.16 | 11.72 | -28.37 | N------OG Thr 256<br>O------NZ Lys 147 | 2.74<br>2.44 | H-don<br>H-acc |
| 3 | Brefeldin A | -10.11 | 6.532 | -17.363 | H------O Arg 89<br>O------OG Thr 256 | 2.06<br>2.78 | H-don<br>H-acc |
| 4 | Emetin | -20.73 | 8.819 | -25.49 | H-------OG Glu 99<br>H-------OG Glu 99 | 1.49<br>1.57 | H-don<br>H-don |
| 5 | Dehydroandrographolid | -13.42 | 6.574 | -23.56 | H-------O Lue 253<br>H-------O Lue 253<br>O-------NZ Lys 261 | 1.81<br>1.71<br>2.76 | H-don<br>H-don<br>H-acc |
| 6 | Harringtonine | -16.42 | 7.146 | -20.45 | H----------O Glu 99<br>O----------N Glu 99<br>O----------OG Thr 256 | 1.76<br>2.95<br>2.82 | H-don<br>H-acc<br>H-acc |
| 7 | Methyllycaconitin | -13.62 | 7.917 | -17.43 | O----------N Lys 96 | 3.52 | H-acc |
| | **SETD7** | | | | | | |
| 1 | Echinomycin | -19.63 | 18.11 | -52.52 | O------NH Asn 282<br>O------NZ Lys 295 | 2.79<br>2.59 | H-acc<br>H-acc |
| 2 | **Streptonigrin** | -27.16 | 21.05 | -71.93 | O-----OG Ser 340<br>O-------NE His 252<br>O------OH Tyr 337<br>O-----OG Ser 340 | 2.75<br>2.80<br>2.67<br>2.75 | H-don<br>H-acc<br>H-acc<br>H-acc |
| 3 | Brefeldin A | -16.66 | 14.88 | -37.69 | H-------O Gly 264<br>O-------N Gly 336 | 1.48<br>2.65 | H-don<br>H-acc |
| 4 | Emetine | -11.62 | 14.04 | -35.15 | H------OD Asp 338 | 1.99 | H-don |

| S.No | Compound | Dock Score (S) | Binding affinity (pki) | Binding energy (kcal/mol) MM-GB/VI | Bonding interactions | Bond length (Å) | Bond type |
|---|---|---|---|---|---|---|---|
|  |  |  |  |  | H--------OD Asp 338 | 1.62 | H-don |
| 5 | Dehydroandrographolid | -20.01 | 19.23 | -47.45 | O--------NZ Lys 294 | 2.58 | H-acc<br>H-acc |
| 6 | Harringtonine | -14.28 | 9.87 | -30.8 | H--------O Gly 336<br>O---------N Gly 336 | 1.73<br>2.73 | H-don<br>H-acc |
| 7 | Methyllycaconitin | -14.16 | 10.02 | -38.23 | O--------NE Trp 352 | 2.68 | H-acc |
| | **NSD3** | | | | | | |
| 1 | Echinomycin | -33.48 | 25.49 | -64.32 | O------NE2 His 1383<br>NH----OG Ser 1406<br>N------OG Ser 1406<br>O-------Zn<br>O----------Zn | 3.4<br>2.9<br>2.7<br>2.4<br>2.2 | H-acc<br>H-acc<br>H-don<br>Ionic<br>Ionic |
| 2 | Streptonigrin | -39.80 | 26.34 | -60.67 | O------NE2 Phe1380<br>O------NE2 His 1408<br>O------Zn | 3.0<br>2.7<br>1.8 | H-acc<br>H-don<br>Ionic |
| 3 | Brefeldin A | -30.48 | 19.24 | -58.12 | O------OG Ser 1406<br>O------Zn<br>O------Zn | 3.1<br>2.2<br>2.1 | H-acc<br>Ionic<br>Ionic |
| 4 | Emetine | -20.65 | 8.45 | -43.22 | O------Zn<br>O------Zn | 2.6<br>2.4 | Ionic<br>Ionic |
| 5 | Dehydroandrographolid | -26.92 | 8.88 | -40.34 | O------Zn<br>O------Zn | 2.1<br>2.1 | Ionic<br>Ionic |
| 6 | Harringtonine | -19.76 | 11.23 | -37.38 | O------Zn<br>O------Zn | 2.08<br>2.2 | Ionic<br>Ionic |
| 7 | Methyllycaconitin | -17.22 | 9.04 | -48.26 | O------Zn<br>O------Zn | 2.04<br>2.07 | Ionic<br>Ionic |
| | **SMYD2** | | | | | | |
| 1 | Echinomycin | -33.39 | 26.84 | -86.4 | O--------NH His 198<br>N--------OH Tyr 258<br>O--------NZ Lys 309 | 3.33<br>2.76<br>2.64 | H-acc<br>H-acc<br>H-acc |

| S.No | Compound | Dock Score (S) | Binding affinity (pki) | Binding energy (kcal/mol) MM-GB/VI | Bonding interactions | Bond length (Å) | Bond type |
|---|---|---|---|---|---|---|---|
|  |  |  |  |  | O--------OH Tyr 344<br>Aren---Aren Tyr 258 | 2.78 | H-acc |
| 2 | Streptonigrin | -16.08 | 18.45 | -33.73 | O------NE Gln 228<br>O------Zn<br>O------Zn | 3.15<br>1.91<br>2.24 | H-acc<br>Ionic<br>Ionic |
| 3 | Brefeldin A | -7.81 | 8.31 | -22.6 | | | |
| 4 | Emetine | -23.82 | 20.24 | -68.45 | H-------OG Thr 185<br>H-------OE Glu 187 | 1.67<br>1.45 | H-don<br>H-don |
| 5 | Dehydroandrographolid | -14.94 | 16.23 | -28.56 | O-------NE His 137 | 2.81 | H-acc |
| 6 | Harringtonine | -15.27 | 18.88 | -42.3 | H------O Tyr 240<br>O------OH Tyr 258<br>O------OG Ser 257<br>O------OH Tyr 250 | 1.49<br>2.78<br>2.92<br>2.78 | H-don<br>H-don<br>H-acc<br>H-acc |
| 7 | Methyllycaconitin | -18.30 | 21.34 | -58.23 | H-------O Leu 191<br>O-------N His 193<br>O-------NE Gln 345 | 1.87<br>2.81<br>3.47 | H-don<br>H-acc<br>H-acc |
| | **MLL1** | | | | | | |
| 1 | Echinomycin | -28.07 | 22.16 | -76.24 | H-------OE Glu 3872<br>O-------N Met 3884<br>O------NH Arg 3886<br>N-------N Lys 3944<br>O-------N Lys 3954 | 2.34<br>2.85<br>2.75<br>2.92<br>2.63 | H-don<br>H-acc<br>H-acc<br>H-acc<br>H-acc |
| 2 | Streptonigrin | -16.45 | 14.54 | -53.72 | O-------------NH Arg 3841<br>O-------OH Tyr 3883 | 2.52<br>2.63 | H-acc<br>H-acc |
| 3 | Brefeldin A | -13.35 | 14.66 | -57.66 | O--------OH Tyr 3883<br>O-------NH Cys 3841<br>O-------N Cys 3882 | 2.82<br>2.69<br>2.74 | H-don<br>H-acc<br>H-acc |
| 4 | Emetine | -19.23 | 17.78 | -50.33 | H-------OE Glu 3872<br>H-------O Cys 3882 | 1.54<br>1.55 | H-don<br>H-acc |

| S.No | Compound | Dock Score (S) | Binding affinity (pki) | Binding energy (kcal/mol) MM-GB/VI | Bonding interactions | Bond length (Å) | Bond type |
|---|---|---|---|---|---|---|---|
|  |  |  |  |  | Cation----aren Tyr 3883 |  |  |
| 5 | Dehydroandrographolid | -14.83 | 14.52 | -54.49 | O--------N Met 3884 | 2.74 | H-acc |
| 6 | Harringtonine | -20.50 | 18.66 | -61.66 | H-------OE Glu 3872<br>H------OD Asp 3876 | 1.63<br>1.65 | H-don<br>H-don |
| 7 | Methyllycaconitin | -18.17 | 16.83 | -58.23 | O------NE Arg 3886 | 2.59 | H-acc |
| | **MLL3** | | | | | | |
| 1 | Echinomycin | -31.97 | 22.32 | -67.54 | O-------N Met 4826<br>H-------O Met 4826<br>N------NH Arg 4828<br>O-------NE Arg 4828<br>O-----NH2 Arg 4828 | 1.87<br>1.90<br>2.91<br>2.57<br>3.41 | H-acc<br>H-acc<br>H-acc<br>H-acc<br>H-acc |
| 2 | Streptonigrin | -22.37 | 21.88 | -66.14 | O-------OH Tyr 4800<br>H-------O Met 4826<br>O-------OH Tyr 4800<br>O-------OH Tyr 4800<br>O-------NE Arg 4828<br>O------NH Arg 4848 | 2.85<br>1.79<br>2.85<br>3.07<br>2.51<br>2.57 | H-don<br>H-don<br>H-acc<br>H-acc<br>H-acc<br>H-acc |
| 3 | Brefeldin A | -15.05 | 16.23 | -54.39 | H-------O Tyr 4886<br>H-------O Tyr 4886<br>O-------NE Arg 4828 | 1.86<br>1.73<br>2.59 | H-don<br>H-don<br>H-acc |
| 4 | Emetine | -26.47 | 9.39 | -27.32 | H-------OE Gln 4781<br>H-------OH Tyr 4886 | 1.52<br>2.11 | H-don<br>H-don |
| 5 | Dehydroandrographolid | -18.19 | 14.23 | -53.65 | H-------O Val 4824<br>O-------OH Tyr 4884 | 1.66<br>2.81 | H-don<br>H-acc |
| 6 | Harringtonine | -19.76 | 17.37 | -58.64 | H-------OE Glu 4814<br>H-------O Met 4827<br>O-------NE Arg 4828<br>O------NH Arg 4828<br>O------N Arg 4828 | 1.47<br>1.78<br>2.82<br>2.70<br>2.88 | H-don<br>H-don<br>H-acc<br>H-acc<br>H-acc |

| S.No | Compound | Dock Score (S) | Binding affinity (pki) | Binding energy (kcal/mol) MM-GB/VI | Bonding interactions | Bond length (Å) | Bond type |
|---|---|---|---|---|---|---|---|
| 7 | Methyllycaconitin | -17.77 | 17.60 | -50.21 | O-------NE Gln 4781<br>O-------N Met 4826 | 2.98<br>3.46 | H-acc<br>H-acc |
| | **MLL4** | | | | | | |
| 1 | Echinomycin | -19.26 | 15.242 | -44.75 | O-------NE Gln 5407<br>N------NE His 5526<br>O-------NE His 5526 | 2.68<br>3.11<br>2.89 | H-acc<br>H-acc<br>H-acc |
| 2 | Streptonigrin | -27.38 | 16.29 | -52.19 | O-------N His 5526<br>O------NE His 5526<br>O-------Zn | 2.86<br>2.90<br>1.86 | H-acc<br>H-acc<br>Ionic |
| 3 | Brefeldin A | -13.30 | 13.07 | -42.52 | H---------O His 5539<br>O---------NE Gln 5407 | 2.03<br>2.79 | H-don<br>H-acc |
| 4 | Emetine | -22.34 | 13.52 | -48.02 | H---------O Arg 5448 | 1.64 | H-don |
| 5 | Dehydroandrographolid | -13.72 | 13.31 | -24.77 | H------OD Asn 5474 | 1.48 | H-don |
| 6 | Harringtonine | -13.31 | 13.46 | -38.366 | | | |
| 7 | Methyllycaconitin | -17.11 | 12.60 | -26.09 | H------OD Asn 5447<br>H------OD Asn 5447 | 1.70<br>1.85 | H-don<br>H-don |
| | **MLL5** | | | | | | |
| 1 | Echinomycin | -18.81 | 13.11 | -50.18 | O-----------N Val 154 | 2.87 | H-acc |
| 2 | Streptonigrin | -22.00 | 18,80 | -65.08 | H----------O Phe 143<br>O---------NZ Lys 73<br>O---------NZ Lys 84<br>O----------N Ala 150<br>O---------NZ Lys 153<br>O---------NZ Lys 153 | 1.55<br>2.74<br>3.00<br>2.95<br>2.33<br>2.51 | H-don<br>H-acc<br>H-acc<br>H-acc<br>H-acc<br>H-acc |
| 3 | Brefeldin A | -16.63 | 11.03 | -42.64 | H---------O Phe 143<br>O---------NZ Lys 73 | 1.61<br>2.49 | H-don<br>H-acc |
| 4 | Emetine | -17.85 | 12.08 | -43.58 | H---------O Cys 108<br>H--------OH Thr 147 | 2.08<br>1.93 | H-don<br>H-don |
| 5 | Dehydroandrographolid | -14.87 | 9.776 | -39.77 | H---------O Phe 145<br>O---------NZ Lys 73 | 1.65<br>2.88 | H-don<br>H-acc |

| S.No | Compound | Dock Score (S) | Binding affinity (pki) | Binding energy (kcal/mol) MM-GB/VI | Bonding interactions | Bond length (Å) | Bond type |
|---|---|---|---|---|---|---|---|
| 6 | Harringtonine | -17.49 | 15.66 | -51.668 | H--------OD Asp 146<br>O---------NH Arg 62<br>O---------NZ Lys 73 | 2.07<br>2.92<br>2.55 | H-don<br>H-acc<br>H-acc |
| 7 | Methyllycaconitin | -10.7 | 14.9 | -43.57 | O---------NZ Lys 153 | 2.77 | H-acc |

**Table S3.** Bonding characterization, dock scores and binding energies of selective seven compounds against various therapeutic targets, H3K9 methyltransferase enzymes

| S.No | Compound | Dock Score (S) | Binding affinity (pki) | Binding energy (kcal/mol) MM-GB/VI | Bonding interactions | Bond length (Å) | Bond type |
|---|---|---|---|---|---|---|---|
| | **Human G9a-like (GLP, also known as EHMT1)** | | | | | | |
| 1 | Echinomycin | -22.83 | 15.57 | -65.12 | H-------OE Glu 1213<br>N-----NH1 Arg 1214<br>Aren--cation Arg 1214 | 1.89<br>2.10 | H-don<br>H-acc |
| 2 | Streptonigrin | -15.27 | 14.60 | -57.37 | H-----OE Glu 1139<br>O------NH Arg 1159<br>O-------NZ Lys 1219<br>O------NH Arg 1226<br>Aren---Aren Phe 1223 | 1.63<br>2.71<br>2.49<br>2.72 | H-don<br>H-acc<br>H-acc<br>H-acc |
| 3 | Brefeldin A | -13.14 | 12.53 | -28.31 | H--------O Met 1105 | 1.85 | H-don |
| 4 | Emetine | -20.26 | 16.57 | -67.82 | H------OD Asp 1147<br>H-------OE Glu 1213<br>O------NH Arg 1214 | 1.48<br>1.39<br>2.83 | H-don<br>H-don<br>H-acc |
| 5 | Dehydroandrographolid | -16.78 | 14.36 | -54.88 | H-------O Asp 1210<br>H------OE Glu 1213 | 1.81<br>1.72 | H-acc<br>H-don |
| 6 | Harringtonine | -17.68 | 13.86 | -49.78 | H------OE Glu 1139<br>O------OG Ser 1141<br>O------NZ Lys 1219<br>Cation-aren Phe 1223 | 1.29<br>2.90<br>2.96 | H-don<br>H-don<br>H-acc |
| 7 | Methyllycaconitin | -20.34 | 17.7 | -59.09 | | | |
| | **Human G9a EHMT2** | | | | | | |
| 1 | Echinomycin | -20.47 | 17.48 | -61.76 | N-----NH Arg 1123<br>O-----NH Arg 1123<br>O-----NH Arg 1123<br>O-----NE Arg 1123 | 2.98<br>2.64<br>2.85<br>2.76 | H-acc<br>H-acc<br>H-acc<br>H-acc |

| S.No | Compound | Dock Score (S) | Binding affinity (pki) | Binding energy (kcal/mol) MM-GB/VI | Bonding interactions | Bond length (Å) | Bond type |
|---|---|---|---|---|---|---|---|
| | | | | | **Cation-ArenArg 1157** | | |
| 2 | Streptonigrin | -19.59 | 16.75 | -65.13 | H-------O Asp 1153<br>O-------NE Arg 1123<br>O------NH Arg 1123<br>O------NH Arg 1123<br>O------N Asp 1153 | 1.68<br>2.41<br>2.71<br>2.77<br>3.39 | H-don<br>H-acc<br>H-acc<br>H-acc<br>H-acc |
| 3 | Brefeldin A | -9.88 | 14.73 | -46.93 | O------N Asp 1153 | 2.53 | H-don |
| 4 | Emetin | -23.20 | 17.73 | -55.45 | H------OD Asp 1088<br>**Aren-cation Arg 1157** | 1.43 | H-don |
| 5 | Dehydroandrographolid | -15.89 | 14.63 | -48.916 | H---------O Tyr 1154 | 1.70 | H-don |
| 6 | Harringtonine | -19.04 | 20.73 | -62.81 | H-------O Tyr 1154<br>H------OD Asp 1156<br>O-------NE Arg 1123<br>O------NH Arg 1123 | 1.64<br>1.48<br>3.01<br>3.28 | H-don<br>H-don<br>H-acc<br>H-acc |
| 7 | Methyllycaconitin | -13.42 | 17.93 | -58.90 | | | |
| | **SUV39H2** | | | | | | |
| 1 | Echinomycin | -19.05 | 20.88 | -48.08 | H--------OD Asp 187<br>O--------OG Ser 56<br>O--------NZ Lys 189 | 1.68<br>2.67<br>2.56 | H-don<br>H-acc<br>H-acc |
| 2 | Streptonigrin | -21.05 | 22.10 | -22.13 | O--------OG Thr 285<br>O--------N Trp 152<br>O--------NE His 216<br>O-------OG Thr 285<br>O-------OG Thr 285<br>O--------NZ Lys 288<br>O---------NZ Lys 288 | 2.78<br>2.54<br>3.21<br>2.78<br>3.20<br>2.53<br>2.57 | H-don<br>H-acc<br>H-acc<br>H-acc<br>H-acc<br>H-acc<br>H-acc |
| 3 | Brefeldin A | -15.57 | 19.39 | -52.39 | H----------O Arg 150 | 1.74 | H-acc |
| 4 | Emetine | -19.19 | 22.35 | -65.90 | O--------NH Arg 145<br>O--------NH Arg 145 | 2.86<br>2.92 | H-acc<br>H-acc |

| S.No | Compound | Dock Score (S) | Binding affinity (pki) | Binding energy (kcal/mol) MM-GB/VI | Bonding interactions | Bond length (Å) | Bond type |
|---|---|---|---|---|---|---|---|
| 5 | Dehydroandrographolid | -14.26 | 17.78 | -21.37 | O--------OG Thr 285 | 2.75 | H-acc |
| 6 | Harringtonine | -16.64 | 20.93 | -30.07 | H--------OGly 149 | 1.77 | H-don |
| 7 | Methyllycaconitin | -18.23 | 22.30 | -15.28 | | | |
| | **SETDB1** | | | | | | |
| 1 | Echinomycin | -23.76 | 18.122 | -64.91 | O--------NE Gln 84<br>O--------NE Trp 169<br>N--------NH Arg 195<br>O--------NH Arg 205 | 2.75<br>2.99<br>2.74<br>2.62 | H-acc<br>H-acc<br>H-acc<br>H-acc |
| 2 | Streptonigrin | -21.98 | 14.919 | -68.92 | O--------OG Thr 204<br>O--------NH Arg 205 | 3.02<br>2.86 | H-acc<br>H-acc |
| 3 | Brefeldin A | -9.33 | 12.58 | -52.39 | H--------OD Asp 81 | 1.68 | H-don |
| 4 | Emetine | -29.66 | 21.13 | -89.13 | H--------ODAsp 110<br>H--------O Asp 110<br>H-------OE2 Glu 197<br>O--------NE Trp 169<br>O--------OG Ser 203 | 2.03<br>1.53<br>1.69<br>3.19<br>3.06 | H-don<br>H-don<br>H-don<br>H-acc<br>H-acc |
| 5 | Dehydroandrographolid | -12.28 | 8.61 | -49.37 | O--------OG Thr 204 | 2.61 | H-don |
| 6 | Harringtonine | -24.16 | 18.47 | -73.76 | H--------O Gly 111<br>O---------OG Ser 139<br>O--------NH Arg 205 | 1.66<br>2.89<br>3.03 | H-don<br>H-acc<br>H-acc |
| 7 | Methyllycaconitin | -19.89 | 17.80 | -62.20 | | | |
| | **SUV39H1** | | | | | | |
| 1 | Echinomycin | -20.32 | 15.4 | -43.28 | H---------OEGlu 75<br>O---------N Gly 43<br>O---------OH Tyr 67 | H-don<br>H-acc<br>H-acc | 2.19<br>2.58<br>2.64 |
| 2 | Streptonigrin | -21.32 | 16.20 | -38.54 | N---------N Gly 43<br>O---------N Gly 43<br>N---------N Gly 43<br>O---------N Gly 43<br>O---------NZ Lys 81 | H-acc<br>H-acc<br>H-acc<br>H-acc<br>H-acc | 2.79<br>2.42<br>3.01<br>2.85<br>2.80 |

| S.No | Compound | Dock Score (S) | Binding affinity (pki) | Binding energy (kcal/mol) MM-GB/VI | Bonding interactions | Bond length (Å) | Bond type |
|---|---|---|---|---|---|---|---|
|  |  |  |  |  | N---------NZ Lys 81<br>aren----aren Trp 64 | H-acc | 2.77 |
| 3 | Brefeldin A | -9.12 | 6.8 | -14.4 | O----------OH Tyr 67<br>O-----------N Gly 43 | H-don<br>H-acc | 2.75<br>2.54 |
| 4 | Emetine | -20.80 | 18.23 | -40.23 | H----------OE Glu 75<br>O----------N Gly 43<br>O----------N Gly 43<br>aren----aren Trp 64 | H-don<br>H-acc<br>H-acc | 1.62<br>2.71<br>2.98 |
| 5 | Dehydroandrographolid | -10.32 | 10.3 | -14.82 | H---------O Glu 46 | H-don | 1.72 |
| 6 | Harringtonine | -13.79 | 9.83 | -21.00 | H--------OE Glu 75<br>O---------N Gly 43<br>O---------N Gly 43 | H-don<br>H-acc<br>H-acc | 1.60<br>2.62<br>3.07 |
| 7 | Methyllycaconitin | -15.23 | 11.4 | -13.65 | H--------OE Glu 75<br>H--------OE Glu 75 | H-don<br>H-acc | 1.87<br>2.27 |
